# The cancer-associated RBM39 bridges the pre-mRNA, U1 and U2 snRNPs to regulate alternative splicing

**DOI:** 10.1101/2022.08.30.505862

**Authors:** Sébastien Campagne, Daniel Jutzi, Florian Malard, Maja Matoga, Ksenija Romane, Miki Feldmuller, Martino Colombo, Marc-David Ruepp, Frédéric H-T. Allain

## Abstract

Pharmacologic depletion of RNA-binding motif 39 (RBM39) using aryl sulfonamides represents a promising anti-cancer therapy. However, its efficiency correlates with the expression level of DCAF15 which acts at the interface between RBM39, the drug and the E3-ubiquitin ligase. Consequently, the identification of alternative approaches to deplete RBM39 independently of DCAF15 is required. Here, we combined transcriptomic analysis, functional assays, and structural biology to elucidate the molecular mechanisms governing RBM39 homeostasis. Our data revealed that RBM39 autoregulates the splicing of its own pre-mRNA by triggering the inclusion of a poison exon. During this process, RBM39 selects the 3’-splice site of the toxic exon, helps the recruitment of U1 snRNP on its weak 5’-splice site and bridges the 3’-splice site recognition machinery. The elucidation of the molecular mechanisms controlling RBM39 homeostasis provides unprecedented insights into alternative 3’-splice site selection and a solid frame to design alternative anti-cancer therapies.

## Introduction

Alternative pre-mRNA splicing is a crucial step of gene expression that not only increases the coding capacity of the genome but also regulates the transcriptional output of genes by affecting mRNA localisation, translation, and decay. Consequently, aberrant pre-mRNA splicing contributes significantly to diseases, in particular to cancer malignancy through regulating the expression of oncogenes and tumour suppressors (Anczuków et al., 2015; Shkreta et al., 2013; Shuai et al., 2019; Srebrow and Kornblihtt, 2006). Understanding how non-physiological splicing patterns drive tumorigenesis and the survival of cancer cells is crucial for the development of novel therapeutic approaches (Bonnal et al., 2020; Lee and Abdel-Wahab, 2016).

The RNA-binding protein called RNA Binding Motif 39 (RBM39) is overexpressed in several types of cancer (Carvalho et al., 2012; Chai et al., 2014; Dai et al., 2016) and is essential for the survival of many cancer cells including Acute Myeloid Leukaemia (AML) cells (Wang et al., 2019). In AML, RBM39 sustains a network of RNA-binding proteins by splicing of their pre-mRNAs and ensures the correct processing of pre-mRNAs encoding Homeobox protein A9 (HOXA9) targets, such as the Polycomb complex protein BMI-1 and the endothelial transcription factor GATA-2, which are required for leukemogenesis (Faber et al., 2009; Katsumura et al., 2016; Yang et al., 2017). Alternatively, RBM39 may also affect blood cell fitness and erythropoiesis by modulating the splicing of VEGF and EPB41 (Dowhan et al., 2005; Huang et al., 2012). As a consequence, a group of anticancer drugs coined the aryl sulfonamides, that specifically induce the targeted degradation of RBM39, trigger the death of cancer cell lines derived from hematopoietic and myeloid lineages (Han et al., 2017; Ting et al., 2019; Uehara et al., 2017) but also cancer stem cells (Chen et al., 2021) and recently, showed exceptional responses in the treatment of high-risk neuroblastoma models (Nijhuis et al., 2022; Singh et al., 2021). The small molecule acts as a molecular glue between RBM39 and the DCAF15-CRL4 E3 ubiquitin ligase to induce proteasome-mediated degradation of RBM39 which in turn leads to cancer cell death (Bussiere et al., 2020; Du et al., 2019; Faust et al., 2020). Furthermore, the targeted degradation of RBM39 generates *bona fide* neoantigens and elicits anti-tumour immunity, augmenting checkpoint immunotherapy (Lu et al., 2021). However, a major caveat of this innovative approach is the dependence on DCAF15, the adaptor protein acting at the interface between the aryl sulfonamides and the E3 ubiquitin ligase activity (Assi et al., 2018; Singh et al., 2021). Hence, the efficiency of this approach correlates with DCAF15 expression levels. Accordingly, the control of RBM39 homeostasis is important for cancer cell survival, and therefore the identification of alternative approaches to lower the intracellular concentration of RBM39 independently of DCAF15 could be beneficial for cancer treatment.

RBM39 is a ubiquitously expressed SR-like protein and is homologous to U2AF2 (Figure S1), the 65 kDa subunit of the U2AF heterodimer (Prigge et al., 2009). The protein consists of an N-terminal RS domain followed by two putative RNA Recognition Motifs (RRM1 and RRM2). A third RRM (RRM3), sometimes referred to as the U2AF2-homology motif (UHM), lost its ability to interact with RNA and has been evolutionarily repurposed to mediate protein-protein interactions with U2AF2 or the U2 snRNP component SF3B1 (Loerch et al., 2014; Stepanyuk et al., 2016). These contacts explain the presence of RBM39 in early spliceosomal complexes A, B and E (Behzadnia et al., 2007; Deckert et al., 2006; Sharma et al., 2008) and indicate how RBM39 could modulate gene expression during pre-mRNA splicing (Huang et al., 2017; Královičová et al., 2018). Furthermore, the RS domains of RBM39 and U2AF2 were shown to promote liquid–liquid phase separation and to favour 3’-splice site recognition (Tari et al., 2019). In contrast to U2AF2, which specifically binds to polypyrimidine tracts (Sickmier et al., 2006), cross-linking and immunoprecipitation (CLIP) experiments suggest that RBM39 has a different RNA-binding selectivity (Mai et al., 2016). However, the RNA-binding specificity of RBM39 remains to be addressed experimentally in order to understand its regulatory role during pre-mRNA splicing.

To decipher the role of RBM39 in RNA metabolism, we combined transcriptomic analysis, cell-based assays and Nuclear Magnetic Resonance (NMR) spectroscopy. Our data revealed that RBM39 actively participates in splice site selection and autoregulates through a negative feedback loop mechanism as RBM39 controls the inclusion of a poison exon in its own pre-mRNA. All three RRM domains of RBM39 are functionally important and independently contact the splicing machinery at both splice sites as well as its pre-mRNA to achieve its autoregulation. In addition, we deciphered the atomic details of RBM39-RNA recognition in this process. Our data bring unprecedented insights into RBM39-dependent 3’-splice site selection which controls the homeostasis of this cancer-associated splicing factor.

## Results

### RBM39 autoregulates its expression by splicing

To investigate the role of RBM39 in RNA metabolism, we performed RNA-Seq in HeLa cells treated with either control (Ctrl KD) or RBM39 siRNAs (RBM39 KD). To discriminate between off-target effects of the siRNAs and genuine RBM39-dependent events, we rescued the RBM39 KD by co-transfecting RNAi-resistant FLAG-RBM39. Western blotting confirmed efficient depletion of the endogenous RBM39, whereas the exogenous rescue protein was moderately overexpressed (Figure S1). Principal component analysis confirmed a strong clustering of the three biological replicates and revealed that the RBM39 KD is the primary source of variation in our data (Figure S1). We then performed differential expression analysis using DESeq2 (Love et al., 2014) and computed meta p-values comparing both Ctrl versus KD and rescue versus KD (Colombo et al., 2017). Using this approach, we found 7’233 mRNAs whose steady-state levels were affected by RBM39 (Figure 1A). Next, we used DEXseq (Anders et al., 2012) to identify 11’134 alternative exon skipping and 7’275 intron retention events that were altered in an RBM39-dependent manner (Figure 1B-C). Among the 1000 most significant events, 73.3% of the transcriptional changes, 82.7% of the exon skipping events and 93.8% of the intron retention events were rescued, corroborating the quality of our RNA-Seq analysis and indicating that the alterations are directly caused by the activity of RBM39, especially those linked to pre-mRNA splicing. Loss of RBM39 induced both up- and downregulation of mRNAs, as well as increased inclusion and skipping of alternative exons. In contrast, more than 80% of the intron retention events showed decreased splicing efficiencies upon RBM39 KD, indicating that RBM39 predominantly enhances the efficiency of constitutive splicing.

**Figure 1.**
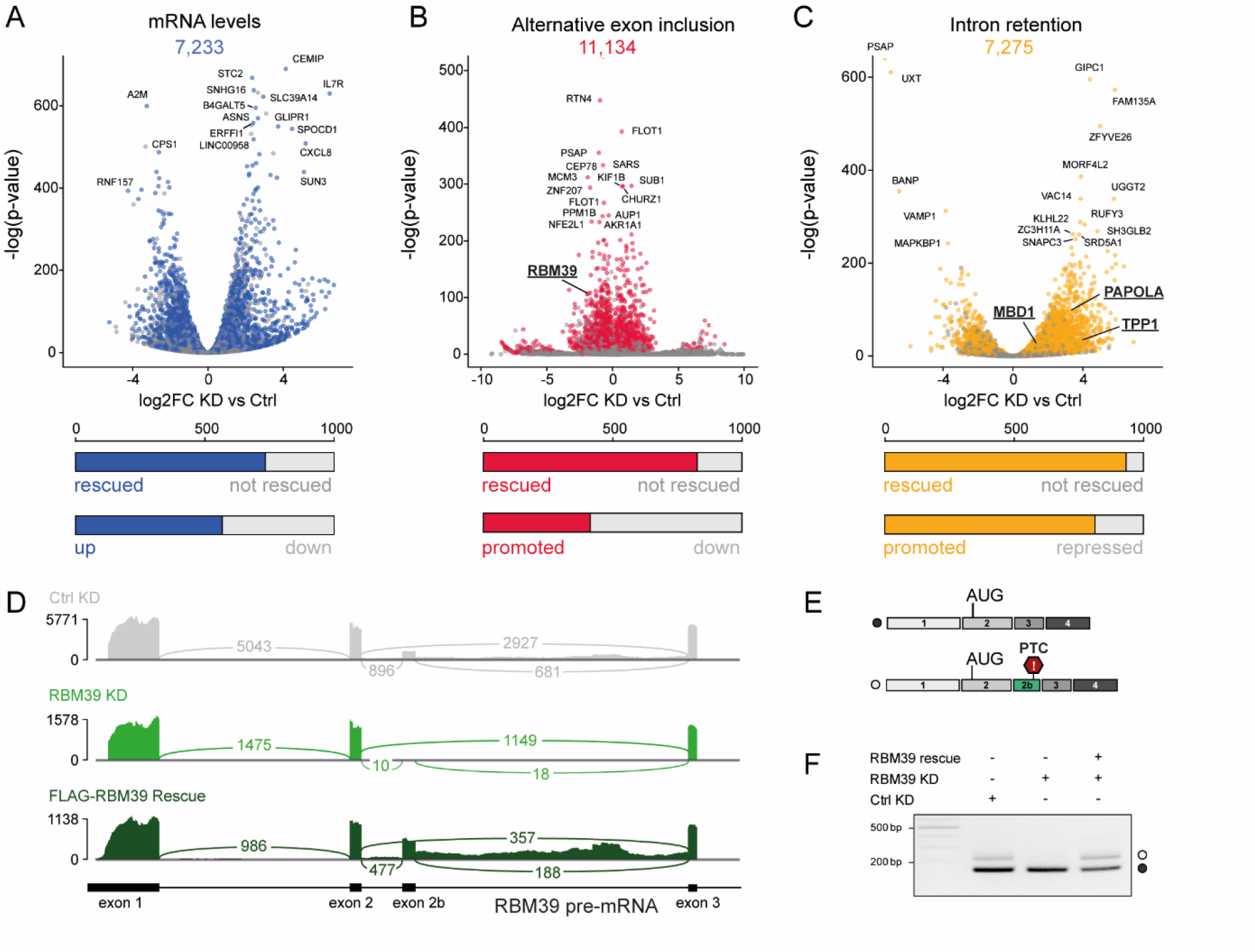
RBM39 controls the inclusion of a poison exon into its own mRNA. A-C) Volcano plots showing up-and downregulation of transcripts (A), alternative exons (B) and retained introns (C) upon RBM39 depletion. Statistically significant alterations (p < 0.05) are highlighted in colour. The bar plots indicate the fraction of rescued events (p < 0.05) and their directionality among the 1000 most significant changes. D) Sashimi plot showing the reads observed for the beginning of the *RBM39* gene upon control knock down (Ctrl KD), RBM39 knock down (RBM39 KD) and FLAG-RBM39 rescue. E) Schematic representation of both RBM39 mRNA isoforms. In the longer isoform, the exon 2b is included and introduces a premature termination codon (PTC). F) RT-PCR validation of the RBM39-dependency of exon 2b inclusion.

The transcriptomic data also revealed that RBM39 promotes the inclusion of a cassette exon (exon 2b) into its own mRNA, which produces an unproductive isoform with a premature termination codon (PTC) and that is predicted to be degraded by the nonsense-mediated decay (NMD) machinery (Schweingruber et al., 2013) (Figures 1D-E). Such PTC-containing exons are sometimes referred to as poison exons. Importantly, this alternative splicing pattern was not caused by the lower levels of RBM39 mRNA in the KD condition, as the expression of FLAG-RBM39 restored the inclusion of exon 2b (Figure 1F). To assess if the PTC-containing isoform is indeed targeted by NMD, we knocked down the central NMD factor UPF1 in HeLa cells using siRNAs. This treatment severely reduced UPF1 mRNA and protein levels and induced a 20-fold upregulation of the endogenous NMD substrate RP9P (Colombo et al., 2017). Total RBM39 mRNA levels were increased about 3-fold upon UPF1 KD, suggesting that in HeLa cells about 66% of the transcripts are degraded under physiological conditions. Using RT-PCR, we could confirm that the PTC-containing mRNA isoform is selectively degraded by the NMD pathway (Figure S1). Altogether, our data show that RBM39 tightly autoregulates its level of expression at the pre-mRNA splicing stage using a negative feedback loop mechanism.

### The three RBM39 RNA recognition motifs contribute to RBM39 function

To support the transcriptomic data, we validated three representative RBM39-dependent intron retention events observed in three pre-mRNA targets (*MBD1, TPP1* and *PAPOLA*, see Figure 1) and investigated the functional contribution of each RRM. Hereto, we rescued RBM39 KD with either FLAG-tagged wild-type RBM39 or mutants lacking one RRM each (Figure 2A-B). All constructs were moderately overexpressed and none of the deletions affected the RBM39 subcellular localisation (Figure 2C). In agreement with our transcriptomic data, RBM39 KD perturbs the splicing of the three introns while the rescue using the wild-type construct either partially or fully restored the splicing defects, indicating that the candidates differ in their sensitivities towards altered RBM39 levels. In particular, TPP1 pre-mRNA splicing may require higher amounts of RBM39 since multiple consecutive introns are retained upon RBM39 depletion (Figure S2). However, for all three intron retention events, the mutant lacking RRM1 (ΔRRM1) did not retain any function, indicating an important role of this domain. Furthermore, the mutants lacking RRM2 (ΔRRM2) or RRM3 (ΔRRM3) only partially rescued the splicing defects compared to the wild-type protein, suggesting that they are also involved in the splicing mechanism (Figure 2D). Interestingly, the functionality profile of the constructs observed for the intron retention events also holds true for alternative splicing of cassette exons, as assessed by the inclusion of the poison exon in the RBM39 mRNA (Figure 2E). In summary, all three RRM domains are important for RBM39-dependent splicing, with a major role played by RBM39 RRM1.

**Figure 2.**
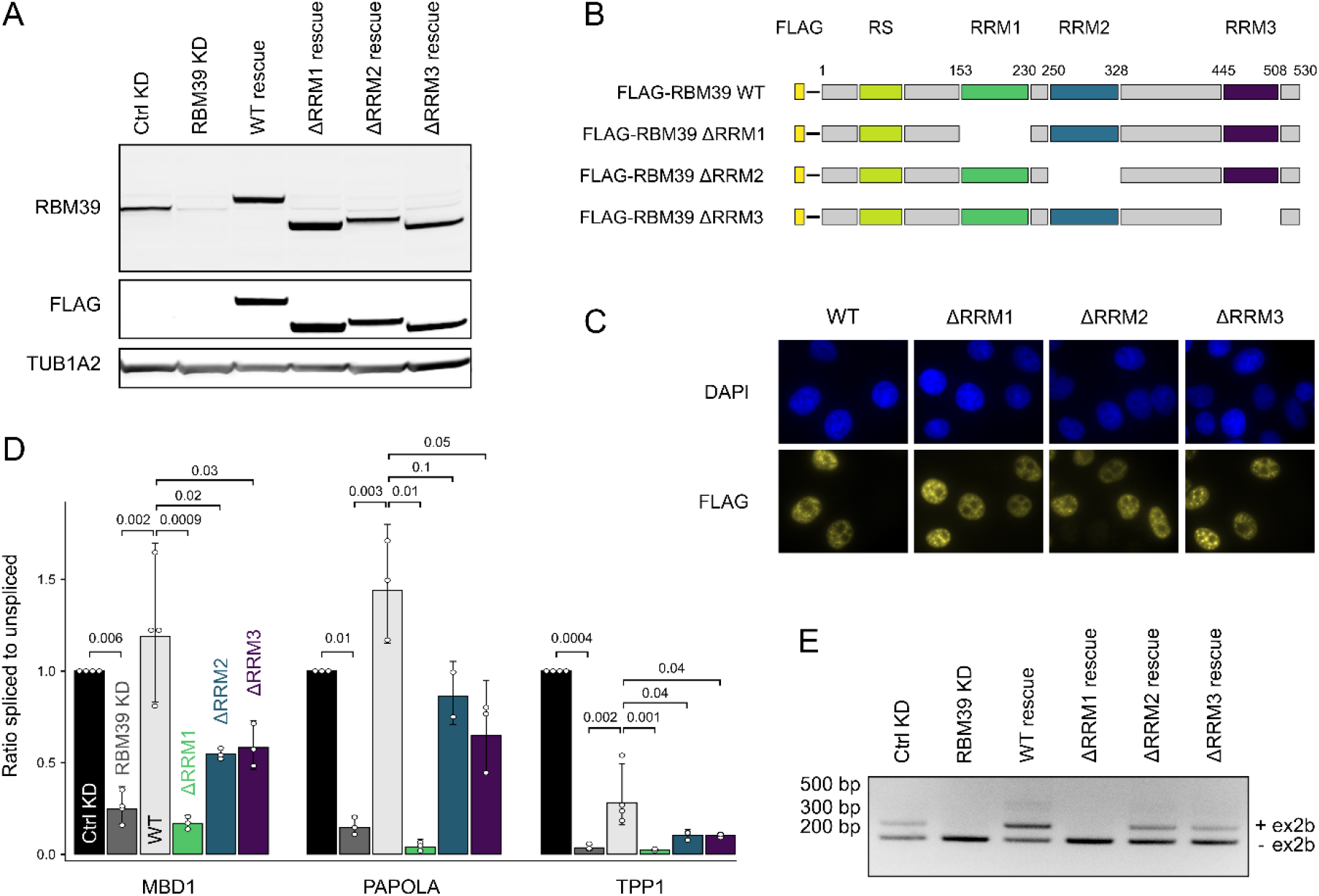
Contributions of the three RBM39 RNA recognition motifs to splicing regulation. A) Western blot analysis of RBM39 levels upon control knockdown (Ctrl KD), RBM39 knockdown (RBM39 KD), FLAG-RBM39 rescue (WT rescue) or rescue performed with RBM39 lacking each one RRM (ΔRRM1, ΔRRM2 and ΔRRM3 rescues). HeLa cell extracts were subjected to SDS–PAGE and Western blotting with anti-RBM39 and anti-FLAG antibodies. Tubulin 1A2 served as a loading control. B) Schematic representation of the FLAG-tagged RBM39 deletion constructs. B) Schematic representation of the FLAG-tagged RBM39 deletion constructs. C) Immunofluorescence analysis of FLAG-RBM39 constructs upon transient expression in HeLa cells. Exogenous RBM39 was visualised using anti-FLAG antibodies and nuclei were counterstained with DAPI. D) RT-qPCR measurements displayed as the ratio of spliced to unspliced isoform for three intron retention events identified by RNA-Seq (MBD1, PAPOLA and TPP1). Average values and standard deviations of three or four biological replicates are shown. P values were computed from log-transformed ratios using two-sided unequal variances Welch’s t-test (Welch, 1947). E) Agarose gel showing the effect of RRM deletion mutants on the splicing of the poison exon of RBM39 mRNA.

### RBM39 bridges early spliceosome components and the pre-mRNA

RBM39, as well as its yeast homolog Rsd-1, were previously identified in early spliceosome complexes (Hegele et al., 2012; Shao et al., 2012). Thus, we investigated potential interactions between RBM39 and the spliceosome components using co-immunoprecipitation in HeLa cell nuclear extracts. In a first attempt, all the U snRNPs were immunoprecipitated with an anti-Y12 antibody (Pisetsky and Lerner, 1982). The isolated complexes contained RBM39 and the U1 snRNP-specific protein U1C (Figure 3A), confirming that RBM39 interacts with the spliceosome. We then precipitated U1 snRNP with an anti-U1A antibody, which again pulled down RBM39 and U1-C (Figure 3A), confirming that RBM39 interacts with U1 snRNP. While this interaction was previously proposed to occur via the RS-domains of RBM39 and U1-70K (Královičová et al., 2018), the association of RBM39 and U1 snRNP was sensitive to RNA digestion, indicating the involvement a protein-RNA contact (Figure 3B). In contrast, the interaction between RBM39 and the U2 snRNP was RNA-independent (Figure 3B), in line with previous observations showing that RBM39 RRM3 contacts either U2 snRNP or U2AF2 through protein-protein interactions (Loerch et al., 2014; Prigge et al., 2009; Stepanyuk et al., 2016). In order to test if the interaction between U1 snRNP and RBM39 could be direct, we *in vitro* reconstituted U1 snRNPs as previously described (Campagne et al., 2019a, 2021) and tested a direct interaction with RBM39 using solution state NMR spectroscopy. As the N-terminal RS domain impaired bacterial expression, we prepared an ILV ^13^C-methyl-labelled RBM39 sample encompassing RBM39 RRM1 and RRM2 (amino acids 143-332, hereafter referred to as RBM39 RRM12). Upon addition of U1 snRNP, the ILV methyl chemical shifts of RBM39 RRM12 experienced changes and more particularly those located in RRM1; consistent with a direct interaction with RRM1 (Figure 3C). The structure of U1 snRNP (Pomeranz Krummel et al., 2009) revealed two protein-free and solvent exposed stem loops (SL3 and SL4) which represent major hubs for the communication with splicing factors (Campagne et al., 2021; Jobbins et al., 2021; Jutzi et al., 2020; Martelly et al., 2021; Sharma et al., 2011). Interestingly, similar ILV methyl chemical shifts changes of RBM39 RRM12 were observed when the protein was titrated with isolated U1 snRNA SL3 (Figure 3C), suggesting that RRM1 binds SL3 when bound to U1 snRNP. According to the RNA dependency of the RBM39-U1 snRNP interaction, we further investigated the interaction between ^15^N-labelled RBM39 RRM12 and *in vitro* transcribed U1 snRNA SL3 using NMR spectroscopy to observe changes in the backbone amides (Figure 3D). As expected, upon addition of U1 snRNA SL3, the amide signals of RBM39 RRM1 experienced fairly large chemical shift perturbations (CSP) confirming that RRM1 is responsible for a direct interaction with U1 snRNP through SL3 (Figure 3D). Using isothermal titration calorimetry (ITC), we determined that RRM1 binds to SL3 with a K_d_ of 15 ± 3 µM (Figure S3).

**Figure 3.**
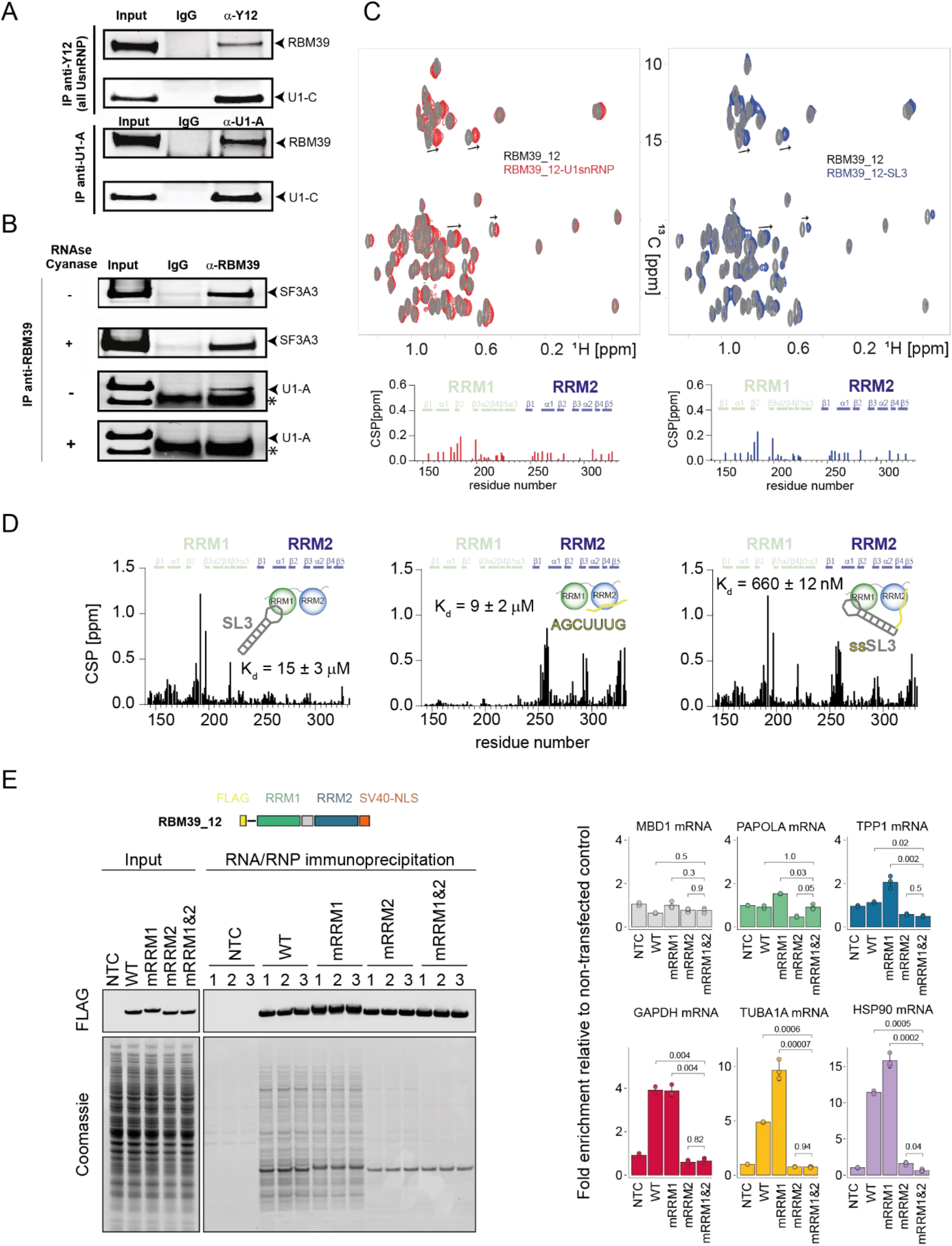
RBM39 interacts with the pre-spliceosome components through protein-protein and protein-RNA interactions. A) Immunoprecipitations performed using an antibody anti-Y12 (top) or anti-U1A (bottom) that pull down all the spliceosome components or U1 snRNP, respectively. The mwestern blots were revealed with anti-RBM39 and anti-U1C antibodies. B) Immunoprecipitations performed using an anti-RBM39 antibody in absence or in presence of RNAse and Cyanase activities and revealed with antibodies against SF3A3 or U1-A. C) Overlay of the 2D ^1^H-^13^C HMQC spectra of RBM39 RRM12 isolated (grey) and after addition of 1 molar equivalent of *in vitro* reconstituted U1 snRNP (red) or purified U1 snRNA SL3 (blue). Bar plots showing the methyl chemical shift perturbations (CSP) observed upon addition of U1 snRNP or U1 snRNA SL3 are displayed. D) Bar plots showing the amide CSPs of RBM39 RRM12 observed upon addition of either U1 snRNA SL3, the ssRNA motif AGCUUUG or the bipartite RNA motifs AGCUUUG-SL3 (ssSL3). Binding affinities measured by ITC are given (Figure S3). E) RNA/RNP immunoprecipitation using FLAG-RBM39 RRM12. On the left side, a scheme of FLAG-RBM39 RRM12 is shown. Below, a western blot shows the presence of the constructs in the input and after immunoprecipitation. The gel below was stained using Coomassie blue. On the right, the presence of immunoprecipitated mRNAs was quantified using RT-qPCR. We detected the three mRNA targets (MBD1, PAOLA and TPP1) of RBM39 and three housekeeping mRNAs (GAPDH, HSP90 and TUB1A).

To identify the main mRNA binding domain of RBM39, we performed RNA immunoprecipitation (RIP) using FLAG tagged wild-type RBM39 or mutants where either RNA-binding by RRM1 or RRM2 or the UHM-ULM interactions of RRM3 are disrupted by point mutations and measured the levels of co-precipitating mRNA by RT-qPCR. All bait proteins were purified with comparable efficiencies and have rich protein interactomes as indicated by Coomassie staining. Compared to a no-transfection control (NTC) condition, the RBM39-regulated mRNAs MBD1, PAPOLA and TPP1 as well as the housekeeping mRNAs GAPDH, HSP90 and TUBA1A were significantly enriched by FLAG-RBM39. However, we did not observe apparent differences between the mutant constructs, suggesting that RBM39 interacts with mRNPs in a redundant manner involving both protein-protein as well as protein-RNA contacts (Figure S3). We therefore repeated the experiment using FLAG-RBM39 constructs encompassing only RRM1 and RRM2 followed by a heterologous SV40 nuclear localisation signal. Under these simplified conditions, the WT and RRM1 mutant proteins retained their rich protein interactomes and robustly enriched the housekeeping mRNAs (Figure 3E). Surprisingly, we did not detect an enrichment of the RBM39-regulated mRNAs with this minimal RBM39 constructs, suggesting that the RS and RRM3 domains are important for substrate specificity. However, the RRM2 and RRM12 mutants lost their ability to bind any mRNA and consequently display strongly reduced protein interactomes, indicating that RRM2 is the main mRNA-binding interface of RBM39.

Using Nuclear Magnetic Resonance (NMR) spectroscopy, we tested the binding of RBM39 RRM12 *in vitro* on different single stranded RNA (ssRNA) motifs that were previously isolated by CLIP (Mai et al., 2016). In line with the RIP experiments, the best CLIP-derived sequences mainly induced amide CSPs on the resonances of RRM2 and the binding became optimal when RRM12 was titrated with a ssRNA having as sequence 5’-AGCUUUG-3’ (Figure 3D). While RRM1 preferentially interacts with RNA stem loops, RRM2 binds 5’-AGCUUUG-3’ with a K_d_ of 9 ± 2 µM (Figure S3). Furthermore, upon addition of a composite RNA containing the ssRNA motif fused to the stem loop 3 (ssSL3, Figure 3D), both RRMs experienced CSPs, showing that both domains directly contact the same RNA molecule and enhance the binding affinity (K_d_ = 660 ± 12 nM; Figure S3). Taken together, our results showed that RBM39 would have the ability to bridge two components of the pre-spliceosome (U1snRNP by RRM1 and U2 snRNP by RRM3) and the pre-mRNA via the central RRM2.

### RBM39 RRM1 recognizes the shape of U1 snRNA stem loop 3

In order to decipher the atomic details of the RBM39 RNA recognition, we studied both RNA-binding domains independently using Nuclear Magnetic Resonance (NMR) spectroscopy. As mentioned above, RRM1 binds preferentially to RNA stem loop structures. Upon addition of U1 snRNA SL3 (hereafter referred as SL3) into a sample of ^15^N-labelled RRM1, we observed amide CSPs on the 2D ^1^H-^15^N HSQC spectra in an unusually large part of the RRM which covers β1 (RNP2), the β1−α1 loop (near A161), the β2-β3 loop (near R192), β3 (RNP2), and the β4-β5 loop (near G220) (Figure 4A-B). On the RNA side, the NMR signals from the loop were the most affected (Figure S4). The solution structure of RBM39 RRM1 bound to SL3 was determined using 2’
s576 NOE-derived distances including 66 intermolecular restraints (Table S1, Video S1 and Figure S4). The ensemble of structures overlaid with a backbone root mean square deviation of 0,68 ± 0,17 Å and revealed that RRM1 interacts with three bases in the RNA loop using its β-sheet surface (A_104_, U_105_ and G_106_). The interactions with U_105_ and G_106_ is only mediated by stacking interactions against the aromatic rings of F156 and Y198 while A_104_ inserts into a cavity along β5 and interacts directly with the side chain of Q159 (Figure 4C-E). The main interaction surface is mediated by the β2-β3 loop that interacts with the major groove of the two base-pairs before the loop and the remaining loop nucleotides (C_101_, A_102_, A_103_ and U_107_). This β2-β3 loop contains four basic side chains (R188, R191, R192 and K194) that establish direct contacts with the phosphate backbone of the loop and with the two GC base pairs at the apical part of the stem (Figure 4F). There are additional interactions with the RNA stem, as the β1−α2 and the β4−β5 loops establish polar interactions with the RNA backbone at the 5’end of the RNA stem. The structure of the protein-RNA complex is in total agreement with the amide CSPs observed upon addition of the SL3 RNA. Its analysis revealed that RRM1 has a poor sequence specificity and rather recognises the shape of this RNA stem loop. The structure of RRM1 bound U1 snRNA stem loop 3 revealed a large positive surface, which perfectly accommodates the RNA loop and its adjacent major groove (Figure S5). Similar CSPs were observed when the protein was titrated with different RNA stem-loops like U1 snRNA SL4 (de Vries et al., 2022) or the stem loop aptamer of RBMY (Skrisovska et al., 2007), confirming a selectivity of RRM1 for stem-loops but with poor sequence specificity (Figure S4). To conclude, the structure of RBM39 RRM1 bound to SL3 explains how RRM1 could associate with U1 snRNP through direct protein-RNA interactions via either SL3 or SL4.

**Figure 4.**
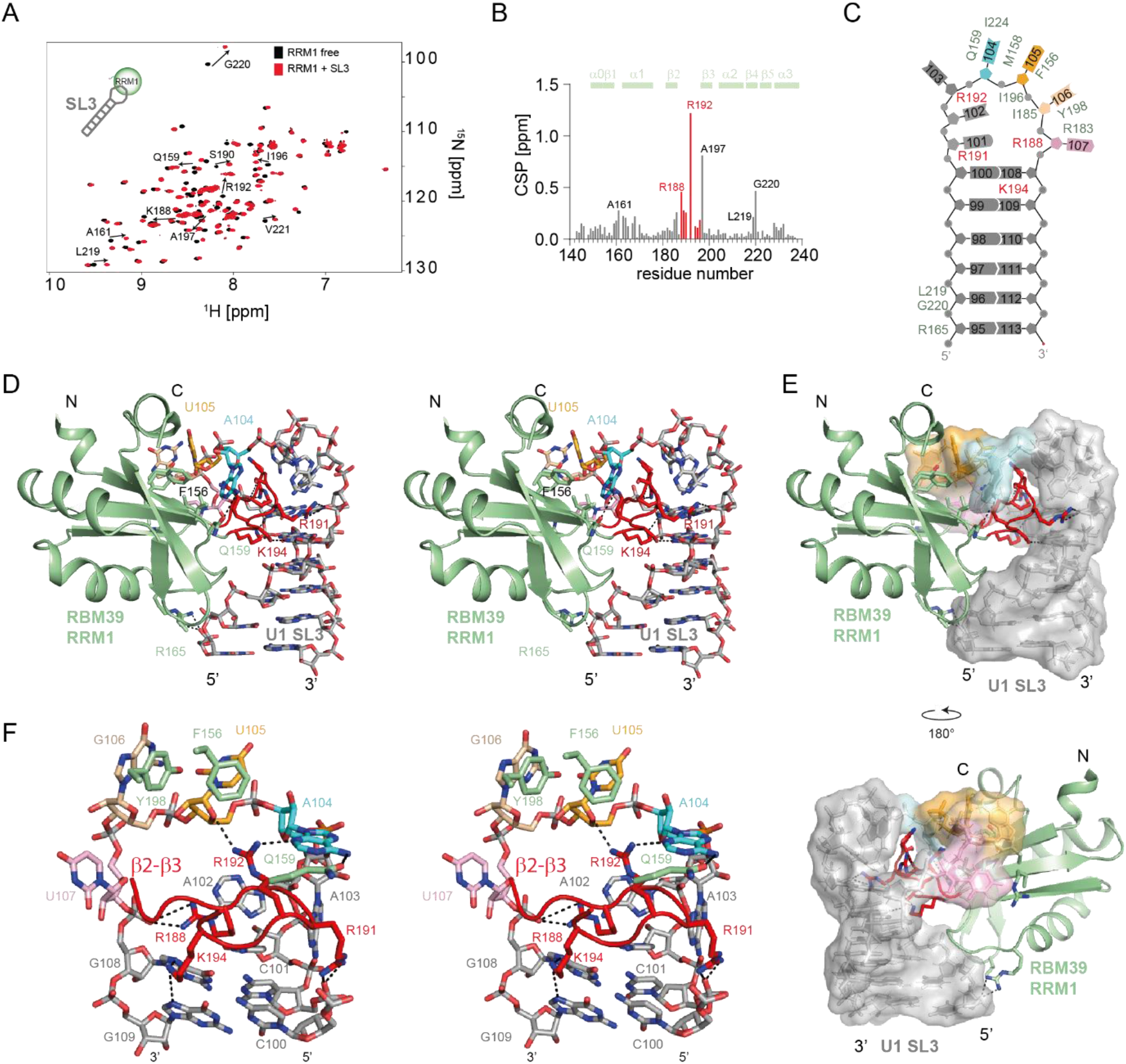
Structural basis for RNA stem loop shape recognition by RBM39. A) Overlay of the 2D ^15^N-^1^H HSQC spectra of ^15^N-labeled RBM39 RRM1 recorded before and after addition of one equimolar amount of U1 snRNA stem loop 3. Spectra corresponding to free and bound protein are coloured in black and red, respectively. B) Normalised amide CSPs are plotted as a function of the sequence of RBM39 RRM1. C) Schematic representation of the protein-RNA contacts observed in the solution structure of RBM39 RRM1 in complex with SL3. D) Stereo view of the solution structure of RBM39 RRM1 in complex with U1 SL3. E) Surface representation. F) Stereo view showing the role of the loop β2-β3 in the recognition of the stem loop shape.

### RBM39 RRM2 binds single stranded RNA motifs using an extended RNA binding interface

Several CLIP consensus motifs (Mai et al., 2016) and derivatives have been assayed for the binding of RBM39 RRM2 and the RNA sequence 5’-AGCUUUG-3’ induced the largest amide CSPs (Figure 5A). Saturation of the CSPs required a slight excess of RNA (1.5-fold excess) and the plots of the amide and carbonyl CSPs as a function of the protein sequence highlighted three main areas of contacts: the loop β1-α1, both β1 (RNP2) and β3 (RNP1) and the C-terminal tail (Figure 5A-B). The solution structure of RRM2 in complex with 5’-AGCUUUG-3’ was determined using 2’
s361 NOE-derived distances including 84 intermolecular distances (Figure 5C, Figure S6, Video S2 and Table S1). The 20 NMR structures overlaid with a backbone RMSD of 0.45 Å and show that RRM2 specifically recognizes two distinct RNA patches (Figure 5C-D). On the β-sheet surface, the 3’-dinucleotide U_6_-G_7_ stacks on the aromatic residues (F295 and Y253) and established direct hydrogen bonds with the backbone of the C-terminal extremity of the protein (Figure 5E). Both preceding nucleotides U_4_ and U_5_ are directly recognised by K322 and R329, respectively, explaining the specificity for the motif UUUG. At the 5’-end, the A_1_-G_2_-C_3_ trinucleotide interacts with the β1-α1 and β2-β3 loops. A key feature of the interaction is the insertion of F259 between A_1_ and G_2_ and the stabilisation of both purines by positively charged amino acids on each side, namely R289 and H258 (Figure 5F). Mutation of F259 into alanine reduced the binding affinity 4-fold, in line with the structure (Figure 5G). Furthermore, the structure revealed a direct hydrogen bond between the O6 atom of G_2_ and the backbone amide group of N260 as well as between the N3 amino of C_3_ and the amide of L256. In agreement, the amide groups of L256 and of N260 are strongly downshifted by the addition of ssRNA. The 5’-terminal A stacks between F259 and R289 and its specific recognition is achieved by the formation of a direct hydrogen bond between the N6 amino and the side chain hydroxyl oxygen of S290. In agreement with the structure, mutations of F259, R289 and H258 to alanine strongly reduced the RNA-binding affinity of RRM2, supporting the important role of the extended interface to achieve high affinity for ssRNA (Figure 5G). Altogether, the solution structure of the protein-RNA complex revealed that RRM2 uses an extended RNA-binding interface that combines the β-sheet surface as well as the α1-β1 and β2-β3 loops.

**Figure 5.**
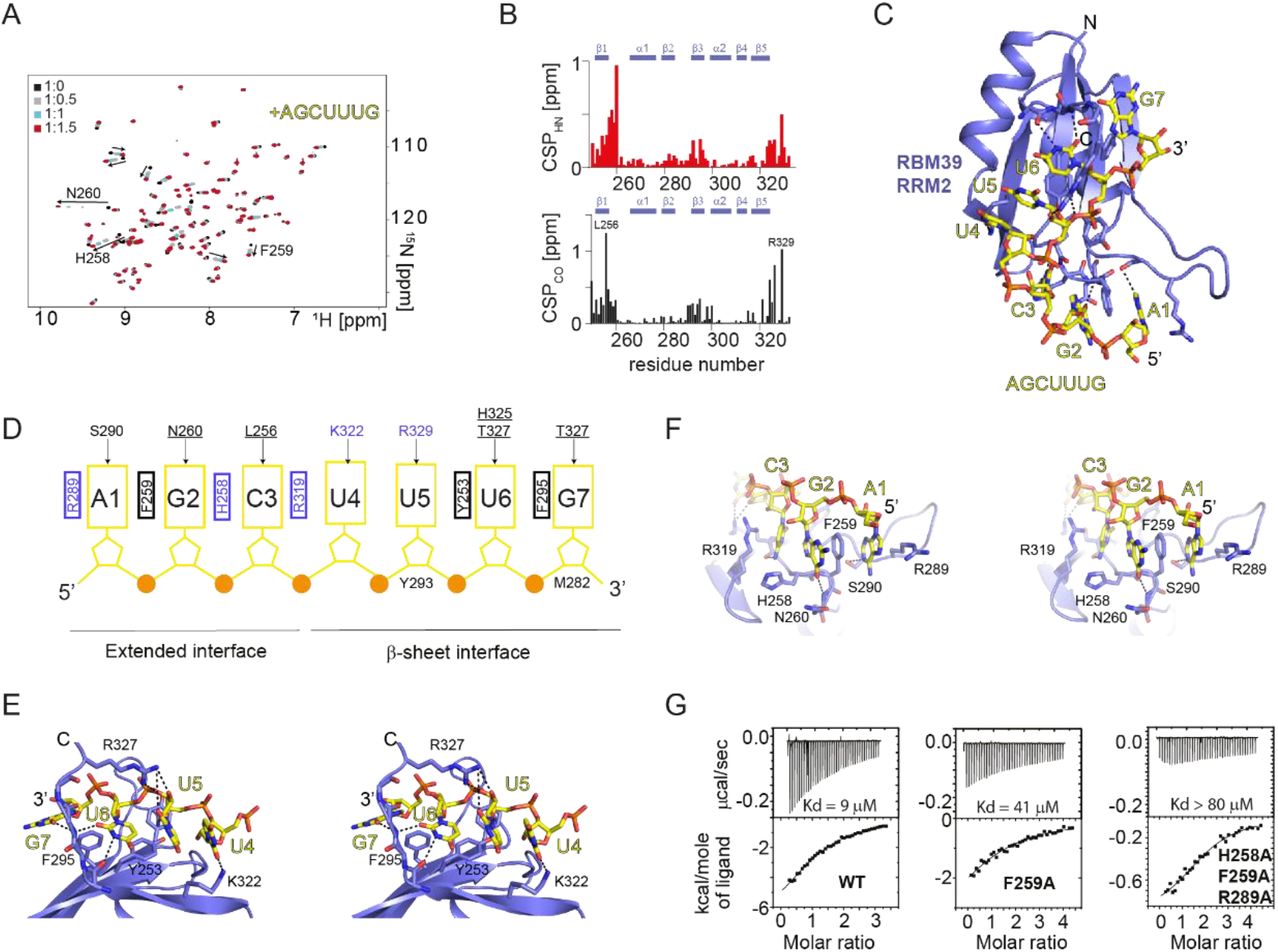
Structural basis for RBM39 single stranded RNA binding activity. A) Overlay of the 2D ^15^N-^1^H HSQC spectra of ^15^N-labeled RBM39 RRM2 recorded upon successive stepwise additions of the 5’-AGCUUUG-3’ ssRNA motif. Spectra are coloured according to the molar ratio protein:RNA (black 1:0; grey 1:0.5; cyan 1:1; red 1:1.5). B) Normalised amide and carbonyl CSPs are plotted as a function of the protein sequence. C) Representation of the lowest energy model of the solution structure of the RBM39 RRM2-AGCUUUG complex. D) Schematic representation of the protein-RNA contacts observed in the solution structure of the RBM39 RRM2-AGCUUUG complex. E) Stereo view of the β-sheet RNA binding interface. F) Stereo view of the extended RNA binding interface involving the loop β1-α1. F) ITC titrations of RBM39 RRM2 variants with 5’-AGCUUUG-3’. The identity of the protein mutants and dissociation constants are given.

### Functional validation of the RBM39-RNA interfaces

In order to test the functional relevance of our protein-RNA complex structures, we mutated key residues at the protein/RNA interfaces and tested the ability of mutants to rescue RBM39 KD on the three intron retention events that we investigated above (Figure 2). We designed a mutant altering RNA-binding by RRM1 on the β-sheet surface (mRRM1.1; F156A/Y198A/R192A), and one mutant altering the contact from the basic residues of the loop β2-β3 (mRRM1.2; R188A/R191A/R192A/K194A). We also prepared an RBM39 mutant altering the RNA-binding interfaces of RRM2 (mRRM2; Y253A/H258A/F259A/F295A), another altering the interaction of RRM3 with U2 snRNP (mRRM3; R494A/F496A) and finally a mutant combining mutations of both RRM2 and RRM3 (mRRM2&3) (Figure 6A). All the proteins expressed at similar levels (Figure 6B) and localised in the nucleus (Figure 6C). Compared to wild-type RBM39, none of the mutants were fully able to rescue the RBM39 KD in all three intron retention events. The mutants having the most severe effects on splicing regulation were mRRM1.1, mRRM1.2 and mRRM2&3 that had an effect comparable to the knock down of the protein (Figure 6D). Similar results were obtained when we tested the mutants on the RBM39 autoregulation splicing event that involves the inclusion of the poison exon 2b (Figure 6E). Altogether, these functional results strongly support our structural findings involving protein-RNA interactions for RBM39 RRM1 and RRM2 (involving non-canonical interactions) and protein-protein interactions for RRM3 (ULM-UHM).

**Figure 6.**
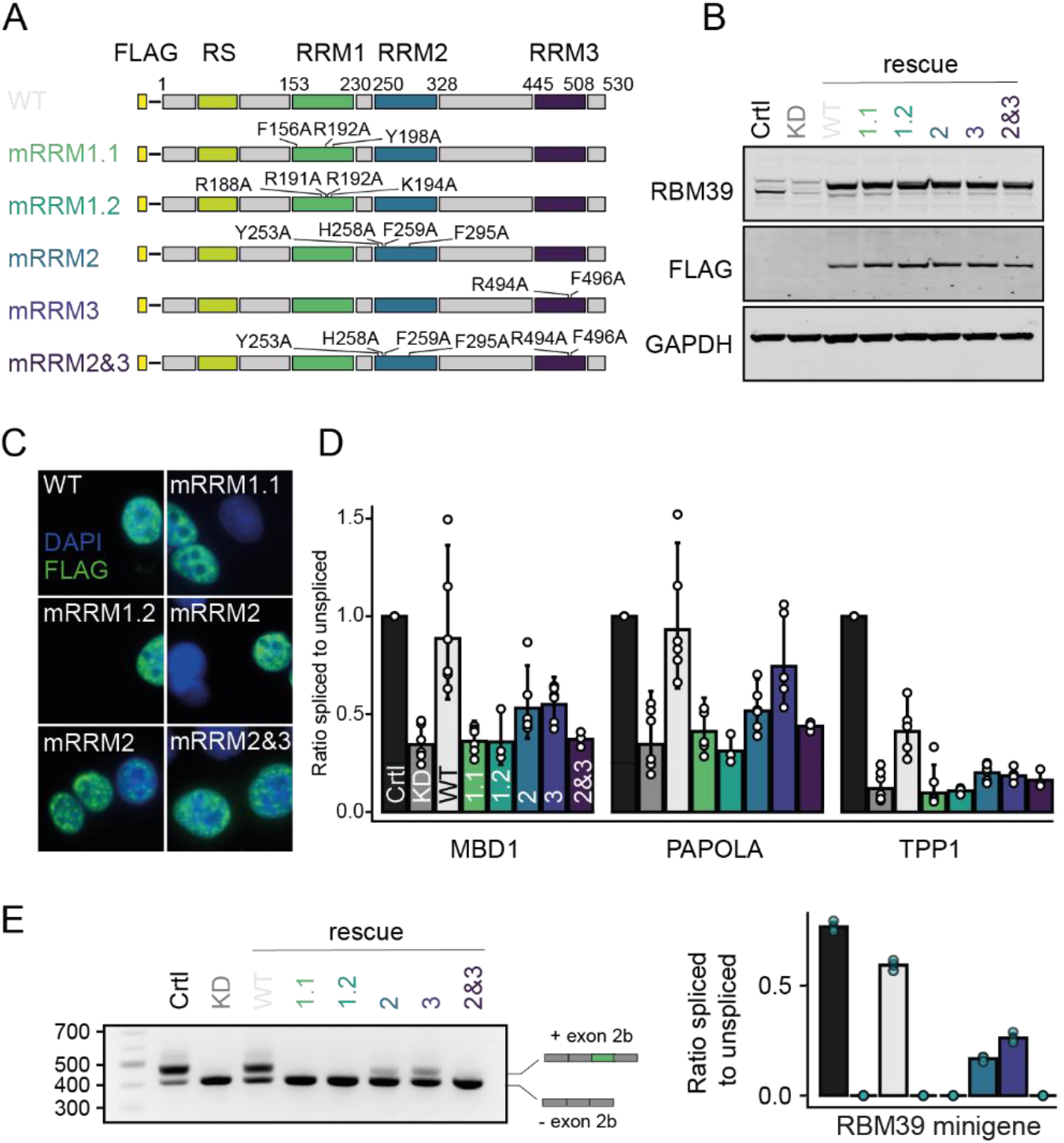
Functional relevance of the RBM39-RNA interfaces in splicing. A) Schematic representation of the RBM39 isoforms used to rescue RBM39 KD. mRRM1.1 is mutated on RRM1 β-sheet surface, mRRM1.2 is mutated on the loop β2-β3, mRRM2 is mutated on the RBM39 RRM2 RNA binding interface and mRRM3 is mutated on its U2 snRNP binding interface. B) Western blot showing the expression of RBM39 variants in eukaryotic cells. C) Immunodetection of the RBM39 variants in cells. All the constructs remained nuclear. D) RT-qPCR measurements displayed as the ratio of spliced to unspliced isoform for three retained introns identified by RNA-Seq (MBD1, PAOLA and TPP1). Average values and standard deviations of three or four biological replicates are shown. P values were computed from log-transformed ratios using two-sided unequal variances Welch’s t-test (Welch, 1947). F) RT-qPCR measurements displayed as the ratio of spliced to unspliced isoform for the poison exon of

### RBM39 autoregulates its splicing through alternative 3’-splice site selection

In order to decipher the molecular mechanisms of RBM39 autoregulation, we investigated how RBM39 promotes poison exon inclusion in a heterologous context. To identify the *cis* RNA elements responsible for RBM39-dependent splicing, we inserted the poison exon and approximately 100 nucleotides of its flanking intronic regions into the β-globin gene between exons 2 and 3. In this heterologous context, the RBM39-dependency of poison exon inclusion was nicely preserved, indicating that the *cis* regulatory elements are located in the poison exon and its flanking regions (Figure S7). Then, we analysed the sequence of the poison exon and identified two potential binding sites for RBM39 RRM2 at key places: CUCUUUG (referred as BS1) and ACCUUUG (referred as BS2) (Figure 7A). While BS1 is located immediately upstream of the AG dinucleotide of the 3’-splice site, BS2 is located within the exon and could potentially form an inhibitory stem loop structure (SLi) by pairing with the 5’-splice site of the poison exon (Figure 7A). To identify the *cis* RNA element responsible for RBM39-dependency, we either deleted BS2 (ΔBS2), substituted the intronic sequence at the 3’-splice site with the β-globin 3’-ss (HBB 3’ss) or specifically mutated BS1 (mutBS1) using our minigene systems (Figure 7A) and evaluated the effect of the mutations on the splicing of the poison exon upon Ctrl or RBM39 KD (Figure 7B). Compared to the endogenous RBM39 mRNA, poison exon inclusion appears to be more efficient in the minigene, likely because the minigene mRNA is less susceptible to nonsense-mediated decay. However, upon RBM39 KD, the percentage of poison exon inclusion in the minigene dropped to 35%. In the context of ΔBS2, RBM39 KD had a moderate but significant effect and reduced poison exon inclusion to 90% (Figure 7B). This decrease in effect size compared to the wild-type minigene could be explained in a scenario where deletion of BS2 disrupts the predicted inhibitory stem loop (SLi) and thus activates poison exon inclusion in an RBM39-independent manner (Figure S7). These results suggest that BS2 is not the primary binding site for RBM39 but represses poison exon inclusion, most likely via sequestration of the weak 5’-ss (Figure 7B). In agreement, the stabilisation of SLi (OPT-SLi) reduced poison exon inclusion (Figure S7). Then, we substituted the upstream region of the poison region by a sequence flanking another exon of the HBB minigene known to be constitutively spliced (Figure 7B and Figure S7). In this genetic context, we lost the RBM39-dependency of the poison exon splicing since no difference was observed between Ctrl or RBM39 KD conditions. Finally, we specifically substituted BS1 by a CCUCCCA motif as in the HBB 3’ss construct (Figure 7B and Figure S7). This subtle change was sufficient to render poison exon inclusion RBM39-independent. In line with our functional data, we could confirm using NMR spectroscopy that RBM39 RRM2 binds strongly to BS1 and BS2 but not to mutBS1 (Figure 7C). Altogether, our data show that the inclusion of the *RBM39* poison exon is controlled by an inhibitory stem loop that may sequester the 5’-splice site and by an RBM39-dependent alternative 3’-splice site selection. In binding to BS1 with its RRM2, RBM39 autoregulates its expression level by enhancing the inclusion of the poison exon 2b.

**Figure 7.**
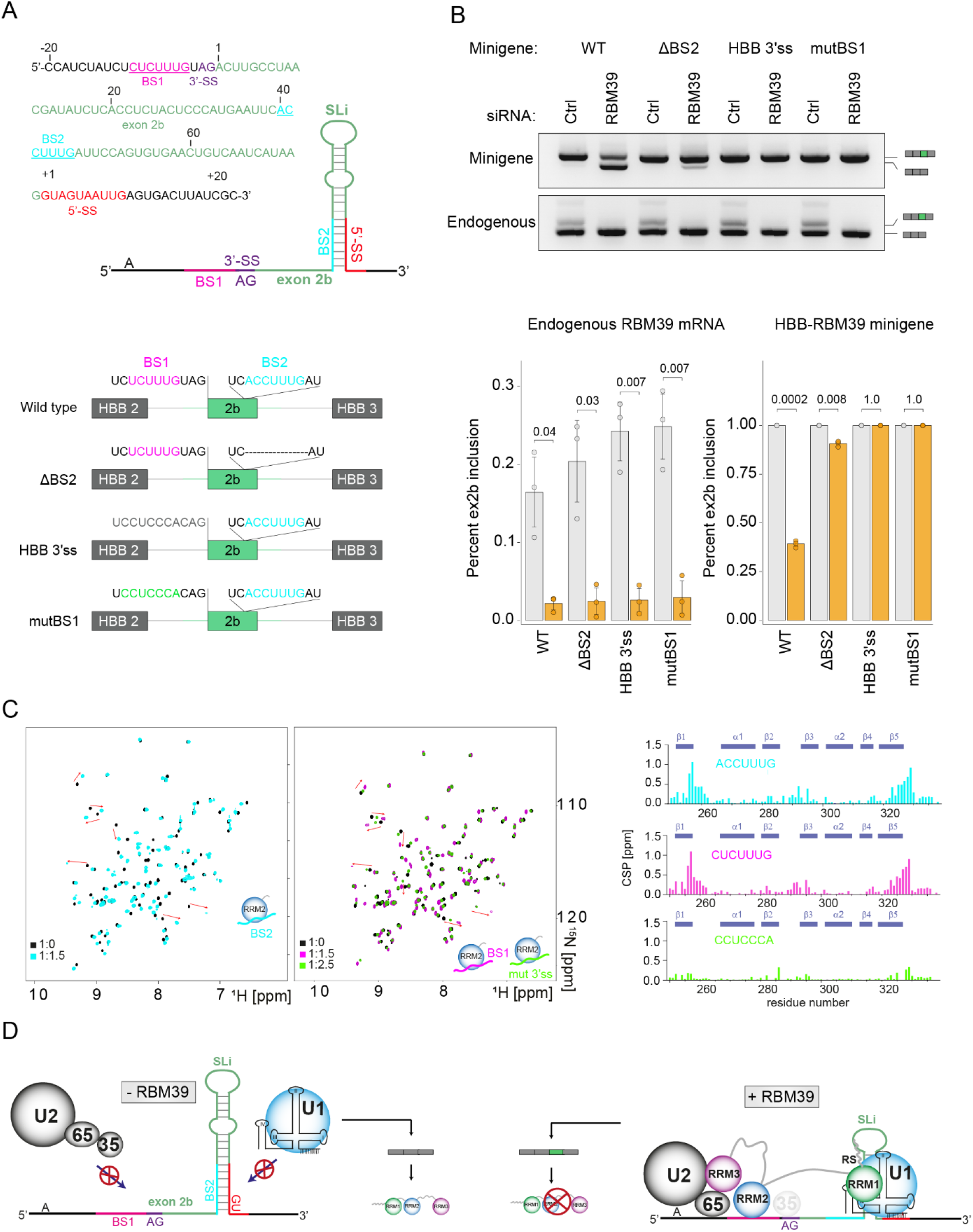
RBM39 controls its homeostasis through alternative 3’-splice site selection. A) Annotated sequence and predicted secondary structures of the *RBM39* exon 2b and its flanking regions. Below, the hybrid minigene containing the exon 2b in the context of the β-globin gene and the different constructs (WT, ΔBS2, HBB 3’ss and mutBS1) are schematically depicted. Sequences are given in Figure S6. B) Percentage of poison exon inclusion was determined by RT-PCR for the different constructs upon Ctlr KD and RBM39 KD. The splicing of the endogenous exon 2b was also monitored in parallel. The percentage of exon 2b inclusion is plotted for each construct in both conditions (Ctlr KD and RBM39 KD). C) Overlay of the 2D ^15^N-^1^H HSQC spectra of RBM39 RRM2 free (black), upon addition of 1.5 equivalents of ACCUUG (blue) or CUCUUUG (pink) or 2.5 equivalents of CCUCCCA. The amide CSPs are plotted as a function of the sequence of RBM39 RRM2. D) Model for poison exon inclusion in the context of RBM39 autoregulation.

## Discussion

### RNA recognition by RBM39 RRM1 and RRM2 are very different

In contrast to its homologs U2AF2 and PUF60, which recognize polypyrimidine stretches using two RRMs in tandem (Figure S1), RBM39 has a different RNA-binding mode and binding specificity. The solution structures of both N-terminal RRMs of RBM39 bound to their respective RNA targets revealed that RRM1 has a strong preference for RNA stem loop structures while RRM2 specifically interacts with 5’-N(G/U)NUUUG-3’ motifs. When compared to the RRMs of FUS and RBMY, which are also specific for RNA stem loop (Jutzi et al., 2020; Loughlin et al., 2019; Skrisovska et al., 2007), the structure of RRM1 bound to SL3 revealed a common strategy used by the three RRMs for the recognition of the stem loop shape. All three RRMs use their β-sheet surface to interact with bases of the loop and insert a loop (β2-β3 in the case of RBM39 and RBMY and β1-α1 in the case of FUS) in the adjacent major groove (Figure S5). This interaction explains the RNA-dependency of the interaction between RBM39 and U1 snRNP that is most likely strengthened by additional RS/RS interactions (Královičová et al., 2018). In contrast, RRM2 anchors the splicing regulator to the pre-mRNPs, as shown by the RIP experiments. The structure of RRM2 bound to its ssRNA target uncovered an extended RNA-binding interface with contacts to seven nucleotides that combines the β-sheet surface and the loops β1-α1 and β2-β3 of the domain. A similarly extended RNA-binding interface was found earlier in Rb-Fox1 RRM, however, the RNA-binding affinity of the Rb-Fox1 RRM for UGCAUGU is 18,000-times stronger (Auweter et al., 2006). This large difference could be explained by the formation of an intramolecular RNA base pair between G_2_ and A_4_ and a larger network of intermolecular hydrogen-bonds in the case of Rb-Fox1 compared to RBM39 RRM2 (Figure S6). According to the structure and the *in vitro* binding assays, RRM2 preferentially binds to the ssRNA motif through its β-sheet face. In contrast, the aryl sulfonamides glue RBM39 and the DCAF15-CLR4 ubiquitin ligase via the helical face of RRM2 (Bussiere et al., 2020; Du et al., 2019; Faust et al., 2020). Since RRM2 uses two opposite faces to contact the RNA and the drug, aryl sulfonamides could also trigger the targeted degradation of the RNA bound state of RBM39. Although the binding affinities of both RRMs for their cognate RNA elements are in the micromolar range, the combination of both RNA elements on the same molecule conferred much higher affinity for RBM39 (20 to 30-fold increase, Figure 3E). This suggests that RBM39, as observed in the case of FUS, could bind RNA targets with a bipartite RNA recognition mode (Loughlin et al., 2019). This property could be used to select bipartite high affinity motifs on pre-mRNA targets, or maybe more importantly, to link two different RNAs molecules; i.e. the pre-mRNA bound by RRM2 and U1 snRNA bound by RRM1.

### RBM39 connects U1, U2 snRNPs and the pre-mRNA

Current models for pre-mRNA splicing regulation by splicing factors mostly propose a recruitment of the splicing machinery to weak splice sites (either 5’ or 3’SS). Typically, SR proteins bind exonic splicing enhancers and recruit U1 snRNP or U2AF1 to the 5’SS or the 3’SS, respectively, using protein-protein interactions (RS domain interactions) (Bourgeois et al., 2004). Other factors like the protein TIA-1 recruits U1 snRNP to the 5’SS by a contact to U1-C again via protein-protein interactions (Forch, 2002). More recently we proposed that FUS recruits U1 snRNP via a direct interaction to U1 snRNA (Jutzi et al., 2020). We also showed that the SR protein SRSF1 can bind U1 snRNA SL3 and many more RNA binding proteins were found to cross-link to U1snRNA (Jobbins et al., 2021). Further, U1 snRNA SL4 can be contacted by splicing factors like PTBP1 (Sharma et al., 2011) as well as the U2 snRNP-specific protein SF3A1, allowing a direct link between U1 and U2 snRNPs (de Vries et al., 2022). Our results extend beyond the above models as we show here that RBM39 could in principle regulate pre-mRNA splicing by simultaneously bridging U1 snRNP (using RRM1), the pre-mRNA (using RRM2) and U2 snRNP (using RRM3). With such a set of interactions, RBM39 could enhance splicing by binding either introns or exons and thus contribute to intron or exon definition, respectively (de Conti et al., 2013). This reflects well the fact that upon RBM39 knock down, both intron retention and cassette-exon inclusion events are perturbed (Figure1) and that RBM39 binding sites both in introns near the 3’SS and in exons near the 5’SS or the 3’SS were observed by CLIP-seq (Mai et al., 2016).

### Molecular mechanisms of RBM39 autoregulation

Among the RBM39-dependent cassette exons, a PTC-containing exon inducing mRNA decay was identified in the *RBM39* gene. Such a negative feedback loop mechanism is commonly observed in autoregulation of key splicing or transcription regulators (Bell et al., 1991; Königs et al., 2020; Leclair et al., 2020; Pervouchine et al., 2019), allowing a precise control of critical gene expression regulator homeostasis. Our experimental data reveal that the inclusion of the poison exon is controlled by two distinct mechanisms. First, our data suggest that RBM39 could stabilise U1 snRNP on a weak 5’-splice site that is sequestered by an inhibitory secondary structure. This strategy is commonly used to regulate alternative splicing of cassette exons such as *SMN2* exon 7 (Singh et al., 2006) or *Map/Tau* exon 10 (Jiang et al., 2000). The stabilisation of U1 snRNP on this weak 5’-splice site could be mediated via the interaction between RRM1 and stem loop 3 of the U1 snRNA; in line with the RNA-dependency of the interaction between U1 snRNP and RBM39. However, this scenario is insufficient to explain the RBM39-dependency of the poison exon, since its inclusion is still sensitive to RBM39 KD in the absence of the inhibitory stem loop (Figure 7). Second, we found that RBM39 controls the inclusion of the poison exon by binding to a short sequence motif located just upstream the AG dinucleotide of the 3’-splice site, in the region −10/−4. The model presented in Figure 7D illustrates the molecular mechanisms of RBM39 autoregulation at the pre-mRNA level. In this process, RBM39 bridges U1 snRNP through its RRM1, the pre-mRNA through RRM2 and the 3’-splice site binding machinery via its UHM-ULM interaction with SF3b155 (Loerch et al., 2014). This mechanism provides an explanation for the modular organisation of the splicing factor and illustrates the concept that the stratification of gene expression control requires multifactorial correlated actions to push the button ‘on’. All three independent domains have a specific task and contribute to RBM39 function in pre-mRNA splicing regulation. The presence of RBM39 at the 3’-splice site most probably remodels the composition of the 3’-splice site recognition machinery and brings several open questions.

Does the UHM interacts with U2 snRNP or U2AF2? Is the function of RBM39 redundant with U2AF1 or U2AF2 or is there a ternary complex possible? Providing answers to these grey areas will provide meaningful insights into the molecular mechanisms of alternative 3’-splice site selection.

### Possibilities for the development of innovative anti-cancer therapy

RBM39 is a major anti-cancer target since its genomic or pharmacological inhibition is detrimental to many cancer cells (Chen et al., 2021; Han et al., 2017; Uehara et al., 2017; Wang et al., 2019). Indeed, following the discovery of the molecular mechanisms triggering the anti-cancer activity of aryl sulfonamides, a clinical evaluation of the combination between chemotherapy and indisulam has been performed (Assi et al., 2018). In AML patients, response to the treatment (30%) correlated with high expression level of DCAF15 and a similar observation was also recently made for the treatment of high-risk glioblastomas using indisulam (Singh et al., 2021). This is in agreement with the mechanism of action of aryl sulfonamides, which act as molecular glue between RBM39 and DCAF15 to induce the degradation of RBM39 and the death of many cancer cells (Bussiere et al., 2020; Du et al., 2019; Faust et al., 2020). This observation suggests that the identification of alternative approaches triggering the degradation of RBM39 independently of DCAF15 could be beneficial for cancer treatments. By deciphering the molecular mechanisms governing the homeostasis of RBM39, we provide important data to manipulate this negative feedback loop mechanism and to trigger the depletion of RBM39 independently of DCAF15. By pushing the splicing equilibrium towards the constitutive inclusion of the poison exon, the expression of RBM39 could be shut down, resulting in cancer cell death as observed with the aryl sulfonamide treatment. The manipulation of the autoregulation mechanism using RNA therapeutics could represent a novel strategy to deplete RBM39 from cancer cells. A similar strategy was recently used to lower the level of *huntingtin* mRNA and develop an innovative therapeutic approach against Huntington’s disease (Bhattacharyya et al., 2021). The rapid expansion of the field of RNA therapeutics correcting splicing has shown the potential of antisense oligonucleotide and small molecule splicing switches to induce targeted splicing correction and has provided novel therapeutic strategies for genetic diseases (Campagne et al., 2019a; Centa et al., 2020; Hua et al., 2011; Lentz et al., 2013; Naryshkin et al., 2014; Palacino et al., 2015; Ursu et al., 2021). In the future, it could also benefit cancer therapy.

## Acknowledgements

This work was supported by the Swiss National Science Foundation (SNF) and the NCCR RNA and Diseases (F.H.-T.A.). This project was sustained by the INSERM Transfert office through the grant COPOC2021 MAT-API-00785-A-01 (S.C.) and by the federal council of La Ligue contre le Cancer through the grant RAB22003GGA (S.C.). This research and related results were made possible through the support of the UK Dementia Research Institute which receives its funding from UK DRI Ltd, funded by the UK Medical Research Council, Alzheimer’s Society and Alzheimer’s Research UK (M.-D.R.), the NOMIS Foundation (M.-D.R.), and the John and Lucille van Geest foundation (M.-D.R.).

## Author contributions

SC, DJ, MDR and FHTA designed the research. SC, MM, FM, KR and MF performed the NMR spectroscopy study. SC solved the structure of the protein/RNA complexes. SC and KR performed the ITC experiments. DJ, MC and MDR performed the transcriptomic analysis and their validation. DJ performed all the cell-based assays. SC, DJ, MDR and FHTA discussed the results and wrote the manuscript.

## Materiel and methods

### Lead contact

Further information and requests for resources and reagents should be directed to and will be fulfilled by the Lead Contact, Frederic Allain.

### Materials availability

All plasmids generated in this study are available upon request from the Lead Contact.

### Data and code availability

The RNA-seq data have been deposited into the GEO database under the accession code GSE202134. The chemical shifts and the structure of the protein-RNA complexes have been deposited in the BMRB (34715, 34673) and PDB (7ZAP, 7Q33).

## EXPERIMENTAL MODEL AND SUBJECT DETAILS

HeLa cells were grown in Dulbecco’s modified Eagle’s medium (DMEM) supplemented with 10% foetal calf serum (FCS), penicillin (100 IU/ml) and streptomycin (100 μg/ml) (DMEM+/+) at 37°C and 5% CO_2_.

## METHOD DETAILS

### Cloning

Plasmids allowing the expression of RBM39 RRM1, RBM39 RRM2 and RBM39 RRM12 were prepared by subcloning the corresponding *E. coli* codon optimized ORF (GeneScript) into pET26bII between the NdeI-XhoI site. RBM39 RRM1 was subcloned using the sc1 and sc2 oligonucleotides; RBM39 RRM2 was subcloned using the sc3 and sc4 oligonucleotides and RBM39 RRM12 was subcloned using the sc1 and sc4 oligonucleotides. Mutagenesis was performed by following the quick-change protocol.

pcDNA3.1-FLAG-GSG15-RBM39 was created by cloning codon optimized RBM39 (GeneArt) into the BamHI and NotI sites of pcDNA3.1-FLAG-GSG15 (gift of Dr. Asimina Gratsou, University of Bern). To generate pcDNA3.1-FLAG-GSG15-RBM39-deltaRRM1 and deltaRRM2, the optimised coding sequences of RBM39 missing amino acids 153 – 230 (RRM1) or amino acids 250 – 330 (RRM2) were ordered from GeneArt, amplified using the primers dj217 and dj218 and inserted into the BamHI and NotI sites of pcDNA3.1-FLAG-GSG15. To create pcDNA3.1-FLAG-GSG15-RBM39-deltaRRM3, the coding region for RBM39 amino acids 1 – 445 was amplified from pcDNA3.1-FLAG-GSG15-RBM39 with dj217 and mdr835 and inserted into the BamHI and NotI sites of pcDNA3.1-FLAG-GSG15.

The point mutations to disrupt RNA-binding and UHM-ULM interactions were introduced by Quick-change mutagenesis of pcDNA3.1-FLAG-GSG15-RBM39. RRM1 was mutated using the primers mdr842 (Y198A), mdr843 (F156A), mdr866 (R192A), whereas RRM2 was mutated using mdr867 (H258A, F259A), mdr864 (Y253A) and mdr865 (F295A). RRM3 was mutated using mdr841 (R494A, F496A).

The pd2-N1-RBM39-minigene was generated as follows: the region spanning exon 1 to 4 of the endogenous RBM39 gene was amplified from human fibroblast gDNA using the primers dj282 and dj283 and inserted into the SalI and NotI sites of pd2-EGFP-N1 (Clonetech). Subsequently, a cryptic splice site in intron 3 was destroyed by QuikChange mutagenesis using the primer mdr848. Pd2-N1-RBM39 deltaBS1 was generated by Quikchange mutagenesis using the primer dj779. To create the pd2-N1-RBM39-optSLi plasmid, the region spanning RBM39 exons 2-3 with the desired point mutations was ordered by gene synthesis and cloned into the EcoRV and Not1 sites of the pd2-EGFP-N1-RBM39 minigene.

To create the RBD-only FLAG-RBM39 constructs for RNA/RNP immunoprecipitation, the RNA-binding region of RBM39 was amplified from the pcDNA3.1-FLAG-GSG15-RBM39 (WT, mutRRM1 and mutRRM2) expression vectors using dj734 and dj735 and cloned into the BamHI and NotI sites pcDNA3.1-FLAG-GSG15. For the RRM1&2 double mutant, the regions encoding RRM1 and RRM2 were first amplified from pcDNA3.1-FLAG-GSG15-RBM39 (mutRRM1 and mutRRM2) using dj734/dj737 (RRM1) and dj736/dj735 (RRM2), and the resulting PCR fragments where then fused by a second round of PCR amplification with dj734/dj735 before cloning into the BamHI and NotI sites of pcDNA3.1-FLAG-GSG15.

To create the pEBFP-c1-HBB minigene with Hind3 and Sal1 restriction sites in intron 2 to facilitate the subsequent introduction of alternative exons, the regions of HBB exon 1 to intron 2 and HBB intron 2 to exon 3 were amplified from fibroblast genomic DNA using the primers dj714, dj715, dj716 and dj717. After fusion of the two fragments by PCR with dj714 and dj717, the HBB gene was inserted into the Bgl2 and Xba1 sites of pEBFP-c1. To generate the pEBFP-c1-HBB-RBM39 WT and deltaBS1 minigene, the region containing RBM39 exon 2b was amplified from pd2-N1-RBM39 minigene WT or deltaBS1 using dj777 and dj778 and cloned into the Hind3 and Sal1 sites of the pEBFP-c1-HBB minigene. To create the pEBFP-c1-HBB RBM39 HBB 3’ss and mutBS1 plasmids, the region spanning RBM39 exon 2b with the desired point mutations was ordered by gene synthesis and cloned into the Hind3 and Sal1 sites of the pEBFP-c1-HBB minigene.

### Protein expression and purification

Expression of RBM39 RRM1, RBM39 RRM2 and RBM39 RRM12 were performed in *E. coli* BL21 DE3. ^15^N and ^13^C uniform isotopic labelling was performed in M9 medium complemented by 1g of ^15^NH_4_Cl and/or 2g of ^13^C glucose. ILV methyl labelling was performed in M9-D_2_O medium in presence of ^15^N-labelled ammonium chloride, unlabelled glucose, 100 mg/L of alpha-ketobutyric acid (methyl-13C, 99%; 3,3-D2, 98%, Cambridge Isotope Laboratory) and 60 mg/L of alpha-ketoisovaleric acid (13C5, 98%; 3-D1, 98%, Cambridge Isotope Laboratory). All the recombinant protein expressions were performed at 37°C during 4 hours in presence of 1 mM IPTG.

Cell lysis was performed using a microfluidizer (3 cycles at 15,000 psi) in buffer A (10 mM Hepes pH7.8, NaCl 1 M, Imidazole 10 mM, β-mercapto-ethanol 2.8 mM) in the presence of DNAse I (10 µg/ml), lysozyme (10 µg/ml) and anti-protease tablets (Roche). The cell lysate was clarified by centrifugation (30,000g, 4°C, 30 minutes) and loaded into an 5ml HisTrap column (Cytiva) previously equilibrated in buffer A. Protein was eluted by a linear gradient of imidazole and dialysed at room temperature during 4 hours in buffer B (10 mM Hepes pH7.5, NaCl 0.25 M, β-mercapto-ethanol 2.8 mM) in presence of thrombin (50 units/10 mg of purified protein; Sigma). The resulting digest was loaded into an 5ml HisTrap column previously equilibrated in buffer B and the flow through was collected, dialysed in buffer C (10 mM sodium phosphate buffer pH 7.0, NaCl 50 mM, DTT 2 mM). The sample was loaded on a 5ml HiTrap SP FF (Cytiva) and eluted with a linear gradient of buffer D (10 mM sodium phosphate buffer pH 7.0, NaCl 1M, DTT 2 mM). RBM39 RRM2 and RBM39 RRM12 were dialysed in buffer E (10 mM sodium phosphate buffer pH 6.8, NaCl 50 mM, DTT 2 mM) and were further purified by size exclusion chromatography (S75, Cytiva) in buffer E. RBM39 RRM1 was dialysed in buffer F (10 mM sodium phosphate buffer pH 5.5, NaCl 50 mM, DTT 2 mM) and was further purified by size exclusion chromatography in buffer F (S75, Cytiva). Recombinant protein purification was monitored by SDS-PAGE.

### RNA production

The RNA U1 snRNA SL3, SL4 and RBMY aptamers were produced by T7-driven *in vitro* transcription using homemade T7 RNA polymerase. Transcriptions were performed using ssDNA templates (Table S2). The transcription mixture was prepared as follow: 30 mM MgCl_2_, 6 mM of each NTP, 4 mM GMP, 2.5 µM dsDNA template and 1.7 µM T7 RNA Polymerase in a transcription buffer containing 40 mM Tris-HCl pH 8.0, 1 mM spermidine, 0.01 % Triton X-100 and 5 mM DTT. 1 U/mL pyrophosphatase was added to reduce magnesium phosphate accumulation. After 4 hours, the reaction was stopped by adding 100 mM EDTA pH 8.0. The mixture was then centrifuged (4,000 x g, 10 minutes) and filtered with 0,22 µm filter. The transcription mixture was then loaded into an preparative anion exchange column mounted on an HPLC system allowing the purification of RNA in denaturing conditions (80°C, 6M urea). The RNA were eluted with a gradient of salt, butanol-extracted, refolded and lyophilised. RNA production and purification were monitored by Urea-PAGE. Short RNA fragments were purchased (Horizon).

### Immunofluorescence

HeLa cells were grown to 80% confluency in 6-well plates and transfected with 300 ng pcDNA3.1-FLAG-RBM39 constructs using Lipofectamine2000 (Invitrogen). On the following day, 540’000 HeLa cells were re-seeded in 8-well chambers (Bioswisstec AG) and fixed after 24 hours with 4% PFA for 30 minutes. After three washes with TBS, the cells were permeabilised and blocked using 1x TBS, 0.5% (v/v) Triton-X-100, 6% BSA at room temperature for 1 hour. Ms anti-FLAG M2 antibodies (Sigma) were diluted 1:200 in TBS+/+ (1x TBS, 0.1% (v/v) Triton-X-100, 6% BSA) and incubated with the cells overnight at 4°C. After 3 × 5 minutes washes with TBS+/+, the secondary antibody (Chicken anti-Mouse AF488, 1:500, Invitrogen) was diluted in TBS+/+ and bound to the primary antibody at 37°C for 1.5 hours followed by incubation at room temperature for 30 minutes. Then, the slides were washed 5 times with 1x TBS and mounted with Vectashield HardSet mounting medium containing DAPI (Vectorlabs). Images were acquired with a non-confocal fluorescence microscope (Leica) using a 60x / 1.4 NA oil immersion lens and the LAS X software (Leica). For printing, brightness and contrast of the pictures were linearly enhanced.

### Immunoprecipitations and western blotting

120 ul Protein G Dynabeads (Life Technologies) per immunoprecipitation were washed three times with TBS supplemented with 0.05% NP-40 (IGEPAL CA-630) and incubated in 600 ul total volume of TBS-0.05% NP-40 with 15 ug of mouse anti U1-70K (Synaptic systems, 203 0111), mouse anti-SmB/B’ (Y12), mouse anti-U1A (SCBT, sc-101149), rabbit anti-RBM39 (Bethyl, A300-291A), mouse IgG (Jackson Immuno Research, 015-000-003), or rabbit IgG (SCBT, sc2027) head over tail for 1.5 hours or supplemented with 1 mg/ml BSA overnight at 4°C. The beads were subsequently washed three times with 1 ml TBS-0.05% NP-40 and resuspended in 600 ul TBS-0.05% NP-40 supplemented with 1 x Halt Protease Inhibitor Cocktail (Thermo Scientific). For RNAse-free IP conditions TBS-0.05% NP-40 was supplemented with 1U/ul NxGen RNAse inhibitor (Lucigen). For nuclease treated samples, the immunoprecipitations were supplemented with 1.5mM MgCl_2_ f.c., 416.6 U/ml Cyanase (Ribosolutions Inc) and 333 ug/ml RNAse A (Sigma). For the RBM39 immunoprecipitations, the reactions were supplemented with 1x Phosphatase inhibitor (Biotool). Hela nuclear extract (Ipracell, CC-01-20-50) was thawed on ice, cleared by centrifugation for 10 minutes at 10’000 x g. 30 ul of cleared nuclear extract was added to the protein G-antibody complexes and incubated for 1.5 hours head of tail at 4°C. Subsequently, the beads were washed three times with TBS-0.1% NP-40, followed by a final wash with 5 minutes head over tail incubation. With the final wash, the beads were transferred to new tubes, wash buffer was removed and the beads were resuspended in 60 μl of 2× LDS-loading buffer, boiled for 10 min at 70°C and loaded on a 4–12% NuPAGE gel. The proteins were transferred on nitrocellulose membranes using the iBlot Gel Transfer Device (Life Technologies). Membranes were blocked with 5% non-fat dry milk in TBS supplemented 0.1% with Tween for one hour and incubated with the primary antibodies rabbit anti-U1C (Bethyl, A303-947A), rabbit anti-RBM39 (Sigma HPA0001591), rabbit anti-SF3b145 (Bethyl, 301-606A), mouse anti-U1A (SCBT, sc-101149), mouse anti-SF3A3 (SCBT, sc-374464) overnight at 4°C. After five washes with TBS-Tween, the membranes were incubated with donkey anti-mouse IRDye800CW (LI-COR Biosciences, 926-32212) or donkey anti-rabbit IRDye800CW (LI-COR Biosciences, 926-32213) in TBS-Tween-Milk for 1.5 hours followed by analysis with the Odyssey Infrared Imaging System (Li-Cor).

### RNA/RNP immunoprecipitation

For the RBM39 FLAG immunoprecipitation, HeLa cells were grown to 50% confluency in 300 cm2 flasks and then transfected with 10 µg of the respective pcDNA3.1 expression vectors using Lipofectamine2000 according to the manufacturer’s protocol. 48 hours post transfection, the cells were harvested with trypsin / EDTA, and the pellets were shock frozen in liquid nitrogen for storage at −80°C until use. To prepare cellular extracts, the pellets were dissolved in 5 mL Gentle Hypotonic Lysis Buffer (10 mM Tris pH 7.2, 10 mM NaCl, 2 mM EDTA, 0.5% Triton-X-100) supplemented with 2x HALT protease inhibitor cocktail (Thermo Scientific, 78429) and 0.5 U/µL RiboLock RNase inhibitor (Thermo Scientific, EO0381) and incubated on ice for 10 minutes. After adjusting the NaCl concentration to 150 mM, the extracts were incubated for another 5 minutes on ice and then cleared by centrifugation at 15,000 x g and 4°C for 15 minutes. Per IP, 100 µL FLAG-M2 matrix (200 µL of a 50% solution) were washed once with matrix preparation solution (50mM Tris pH 7.5, 150mM NaCl) once with 0.2 M Glycine pH 3.5 (to remove unconjugated FLAG antibodies) and again twice in matrix preparation solution. To keep protein and RNA input fractions, 50 μl of the cleared extracts were boiled with 2x LDS loading buffer and 200 μl were transferred to 1 ml TRIzol (Invitrogen, 15596018) supplemented with 0.14 M β-mercaptoethanol (Applichem, A1108). Three times 1.5 mL of the extracts were then distributed to Eppendorf tubes containing FLAG-M2 matrix and incubated on a rotary wheel for 1.5 hours at 4°C. After washing the beads 5 times with HEPES NET-2 buffer (50 mM HEPES pH 7.3, 150 mM NaCl, 0.1 % Triton-X-100), 1/5 of the beads was transferred into a separate Eppendorf tube and boiled in 2x LDS loading buffer, whereas the remaining beads were transferred to 1 ml TriZOL supplemented with 0.14 M β-mercaptoethanol. Proteins were analysed using 4-12% Bis-Tris polyacrylamide gels in MOPS buffer followed by Coomassie staining or western blot using the following antibodies: Mouse anti-FLAG M2 (Sigma, 1:10,000). RNA was isolated from TriReagent according to the manufacturer’s protocol and then reverse transcribed using the high capacity RNA-to-cDNA kit (Applied Biosystems, 4387406) and analysed by RT-qPCR.

### Isothermal titration calorimetry

ITC experiments were performed on a VP–ITC instrument (Microcal). Both partners were prepared in the NMR buffers (buffer E for RBM39 RRM2 and RBM39 RRM12 or buffer F for RBM39 RRM1): the protein (250 µM for RBM39 RRM1 and RBM39 RRM2; 90 µM for RBM39 RRM12) was injected into a solution of ssRNA (10 µM for SL3 and AGCUUUG; 7 µM for the composite RNA motif ssSL3) by 40 injections of 6 µl every 350 s at 35 °C (for RBM39 RRM12) or at 40°C (for RBM39 RRM1 and RBM39 RRM12). Raw data were integrated, normalized for the molar concentration and analysed using Origin 7.0 according to a 1:1 binding model.

### RBM39 knockdown and RNA-Sequencing

HeLa cells, in 6-well plates at 80% confluency were transfected using Lipofectamine 2000 with either 120 pmol of Control siRNA (5’-AGGUAGUGUAAUCGCCUUGdTdT-3’) and 300 ng of pcDNA3.1, 60 pmol RBM 3’UTR (5’-GAGAAUUCAUCUUGAGUUAdTdT-3’), 60 pmol RBM CDS2 siRNA (5’-GGAAAGAGAUGCAAGGACAdTdT-3’) and 300 ng of pcDNA3.1, or 60 pmol RBM 3’UTR, 60 pmol RBM CDS2 siRNA and 300 ng of pcDNA3-FLAG-RBM39 using Lipofectamine 2000. 24 hours post transfection the cells were split and re-transfected the next day with 160 pmol of siRNAs and 300 ng of pcDNA3 expression constructs. The cells were expanded the next day and harvested 5 days post transfection in Tri-Reagent for RNA extraction or RIPA Lysis and extraction buffer (Thermo Scientific) for immunoblot analysis. 1.5 × 10^5^ cell equivalents were loaded on a 4-12% Bolt Bis-Tris protein gel and subjected to western blotting. Knockdown and rescue were analysed using rabbit anti-RBM39 (Bethyl, A300-291A, 1:10,000), mouse anti-FLAG (Sigma-Aldrich, F3165) and rabbit anti-Actin (Sigma-Aldrich, A5060, 1:2,000). RNA from each condition in triplicates was processed at the genomics core facility of the University of Bern using the Illumina TruSeq Stranded mRNA Library Prep Kit and sequenced on an Illumina HiSeq3000 platform using in 2 × 150 bp paired-end sequencing cycles.

To assess the ability of RBM39 mutants to promote pre-mRNA splicing, we rescued siRNA-mediated knockdown of endogenous RBM39 by co-transfection of 300 ng RNAi-resistant pcDNA3-FLAG-RBM39 constructs using the protocol described above. To study the autoregulation of RBM39, 100 ng pd2-N1-RBM39-minigene were included in the second transfection at day 3. UPF1 knockdown experiments were performed according to the same timeline employing either 120 pmol Ctrl siRNA or 120 pmol UPF1 siRNA (5’-GAUGCAGUUCCGCUCCAUUdTdT-3’) along with 300 ng of empty pcDNA3 vector. Western blots were probed with rabbit anti-RBM39 (Bethyl, A300-291A, 1:10’000), mouse anti-FLAG (Sigma-Aldrich, F3165, 1:2’000), goat anti-UPF1 (Bethyl Laboratories, A300-038A, 1: 1,000) and mouse anti-TUB1A2 (Sigma Aldrich, T9028, 1:5’000) to assess knockdown and rescue efficiencies. RNA isolated from TriReagent was DNase treated using the Turbo DNA-free kit (Ambion) followed by reverse transcription using the high-capacity RNA-to-cDNA kit (Applied Biosystems). RT-PCR was carried out for 25 cycles using 2x KAPA Taq ready-mix (Kapa Biosystems) in 50 µl reactions containing 80 ng cDNA and 400 nM primers each, following the manufacturer’s manual. qPCR was performed using the 2x MESA Green qPCR Master Mix Plus (Eurogentec) for SYBR assays in 50 µl reactions containing 32 ng cDNA at a primer concentration of 600 nM each, according to the manufacturer’s instructions. Pipetting was performed using a QIagility pipetting robot (Qiagen) and a Rotor-Gene 6000 Q cycler (Qiagen) was used for PCR amplification and fluorescence monitoring. Primer sequences are listed in Table S2. Relative quantifications of transcripts were carried out using the ΔΔCt method(Livak and Schmittgen, 2001) and statistical significances were assessed on non-transformed ΔΔCt values using the Welch’s test (Welch, 1947) to account for unequal variance between the conditions.

### Transcriptomic analysis

RNA-seq data has been analysed for transcriptional and isoform-level changes. Common to both analyses is the mapping to the human genome GRCh38 by means of STAR aligner version 2.5.2a (Dobin et al., 2013). Annotation indexes were built based on Ensembl GTF files (release 84). Gene-level counts were computed with featureCounts (Liao et al., 2014) program of the Subread package (version 1.4.6). Differential gene expression analysis was carried out with default options with DESeq2 version 1.6.3 (Love et al., 2014). The differential analyses have been carried out on two comparisons: KD vs control and rescue vs KD. A meta-analysis of p-values of the two comparisons determined the list of all significant events. Alternative exon usage was instead identified by using DEXseq (Anders et al., 2012) on BAM files. Ensembl isoforms were filtered for main transcripts as classified in the APPRIS database (Rodriguez et al., 2018). Finally, intron retention was quantified with custom python scripts based on PySam package (Reber et al., 2016). Briefly, a ratio between spliced and unspliced reads was calculated and the significant differences between controls, KD and rescue conditions were assessed using a modified DESeq2 run (https://github.com/Martombo/SpliceRatio).

### In vitro reconstitution of U1 snRNP

The preparation of the U1 snRNP components and the particle assembly was performed as previously described (Campagne et al., 2021).

### NMR spectroscopy

All the NMR spectroscopy measurements were performed either at 308K (for RBM39 RRM2) or at 313K (for RBM39 RRM1 and RBM39 RRM12) using Bruker AVIII 500 MHz, AVIII 600 MHz, AVIII 700 MHz and Avance 900 MHz spectrometers. The data were processed with Topspin3.2 (Bruker) and analysed with CARA (Keller, 2004). Sequence specific backbone and side chain assignments were achieved using the classical approach (Sattler, 1999). All NOE spectroscopy (NOESY) experiments were recorded with a mixing time of 80 ms to avoid spin diffusion. RNA assignment was performed using 2D ^1^H-^1^H homonuclear TOCSY and NOESY for the ssRNA target and U1 SL3. U1 SL3 was also produced ^15^N/^13^C-labelled and assigned by combining 3D HCCH-TOCSY and 3D ^1^H-^13^C HSQC NOESY. Identification of intermolecular NOEs was achieved by comparing 2D F2f ^1^H-^1^H NOESY and 2D F1fF2f ^1^H-^1^H NOESY recorded with various mixing times ranging between 60 ms to 150 ms (Campagne et al., 2019b) for both protein-RNA complexes. For the complex RBM39 RRM1-U1SL3, we also used 3D ^13^C-(F1 edited, F3 filtered) NOESY HSQC (Zwahlen et al., 1997) using RBM39 RRM1 ^15^N-^13^C labelled and U1SL3 unlabelled.

### Structure calculation

To solve the structures of the RNA bound states of RBM39 RRM1 and RBM39 RRM2, the resonance assignments of the bound proteins were used for peak picking and automatic NOE assignment in the 3D NOESY spectra using UNIO–ATNOS–CANDID (Herrmann et al., 2002) in combination with structure calculations using CYANA. In addition, dihedral angle constraints were derived with TALOS+ using backbone chemical shifts as input (Shen et al., 2009) and hydrogen bonds were identified based on reduced amide exchange rates in D_2_O. RNA intramolecular NOEs and protein-RNA intermolecular NOEs were picked manually. The resulting peak-lists including intramolecular and intermolecular NOEs, the hydrogen bonds restraints and backbone dihedral angle restraints were combined to calculate the initial structures of the protein-RNA complex using the CYANA NOEASSIGN module (Güntert and Buchner, 2015). The 50 lowest energy structures were refined in cartesian space using the SANDER module of AMBER20 (Case et al., 2005). Analysis of the refined structures was performed using AMBER20 scripts and PROCHECK (Laskowski et al., 1996).

