## Supplementray data for "The cancer-associated RBM39 bridges the pre-mRNA, U1 and U2 snRNPs to regulate alternative splicing"

[1]ETH Zurich, Department of Biology, Institute of Biochemistry, 8093 Zurich, Switzerland; [2]United Kingdom Dementia Research Institute Centre, Institute of Psychiatry, Psychology and Neuroscience, King's College London, Maurice Wohl Clinical Neuroscience Institute, London SE5 9RT, United Kingdom; [3]University of Bordeaux, Inserm U1212, CNRS UMR5320, ARNA Laboratory, 33077 Bordeaux, France; [4]University of Bern, Department of Chemistry and Biochemistry, 3012 Bern, Switzerland; [5]Celgene Institute of Translational Research (CITRE), 41092 Seville, Spain.

SC and DJ contributed equally to this work.

This supplementary file is composed of:

**7 supplementary figures:**

- Figure S1, related to Figure 1. RBM39 is homologous to U2AF2 and PUF60 and it promotes the inclusion of a poison exon in own pre-mRNA.
- Figure S2, related to Figure 2. RBM39-dependent intron retention events.
- Figure S3, related to Figure 3. ITC measurements and RNA immunoprecipitation using full-length RBM39
- Figure S4, related to Figure 4. RNA spectra, identification of intermolecular NOEs and solution structure of the RBM39 RRM1-U1 SL3 complex.
- Figure S5, related to Figure 4. Structural comparison between RBM39, RBMY and FUS RRM interactions with RNA stem loops.
- Figure S6, related to Figure 5. Identification of intermolecular NOEs, solution structure of the RBM39 RRM2-AGCUUUG complex and comparison with Fox-1 RRM – RNA interaction.
- Figure S7, related to Figure 7. Identification of the cis RNA element responsive for the RBM39 dependency of the poison exon inclusion.

**2 supplementary tables:**

- Table S1 | NMR statistics of the RBM39 RRM1-U1SL3 and RBM39 RRM2-AGCUUUG complexes
- Table S2 | Oligonucleotides list

**2 supplementary videos:**

- Video S1, related to Figure 4. Solution structure of RBM39 RRM1 bound to U1 snRNA stem loop 3.
- Video S2, related to Figure 5. Solution structure of RBM39 RRM2 bound to the ssRNA motif AGCUUUG.

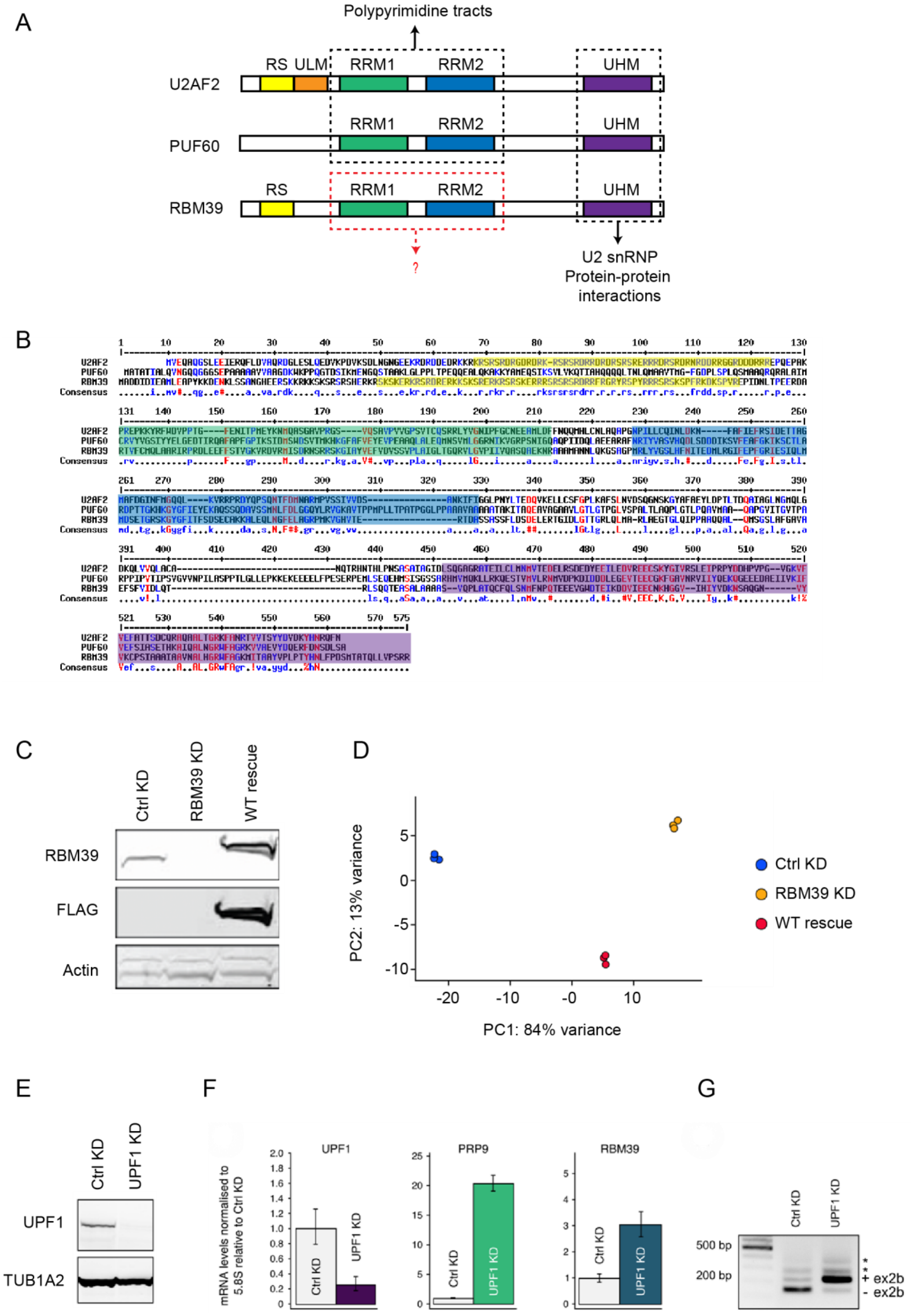


**Figure S1, related to Figure 1. RBM39 is homologous to U2AF2 and PUF60 and it promotes the inclusion of a poison exon in own pre-mRNA.** A) Domain organisation of RBM39, U2AF2 and

PUF60. B) Sequence alignment of RBM39, U2AF2 and PUF60. C) Western blot analysis showing the efficiency of RBM39 siRNA knock down and its rescue with FLAG-RBM39. D) Principal component analysis of the 9 samples that were analysed by RNA-Seq. E) Western blot showing the knock down of UPF1. F) Bar plots showing the mRNA abundance of UPF1, PRP9 and RBM39 in wild type conditions and after UPF1 knock down. G) Agarose gel showing the accumulation of the RBM39 non-productive mRNA isoform in the UPF1 knock down conditions (UPF1 KD). In control conditions (Ctrl KD), the RBM39 non-productive mRNA isoform is degraded via non-mediated decay pathway.

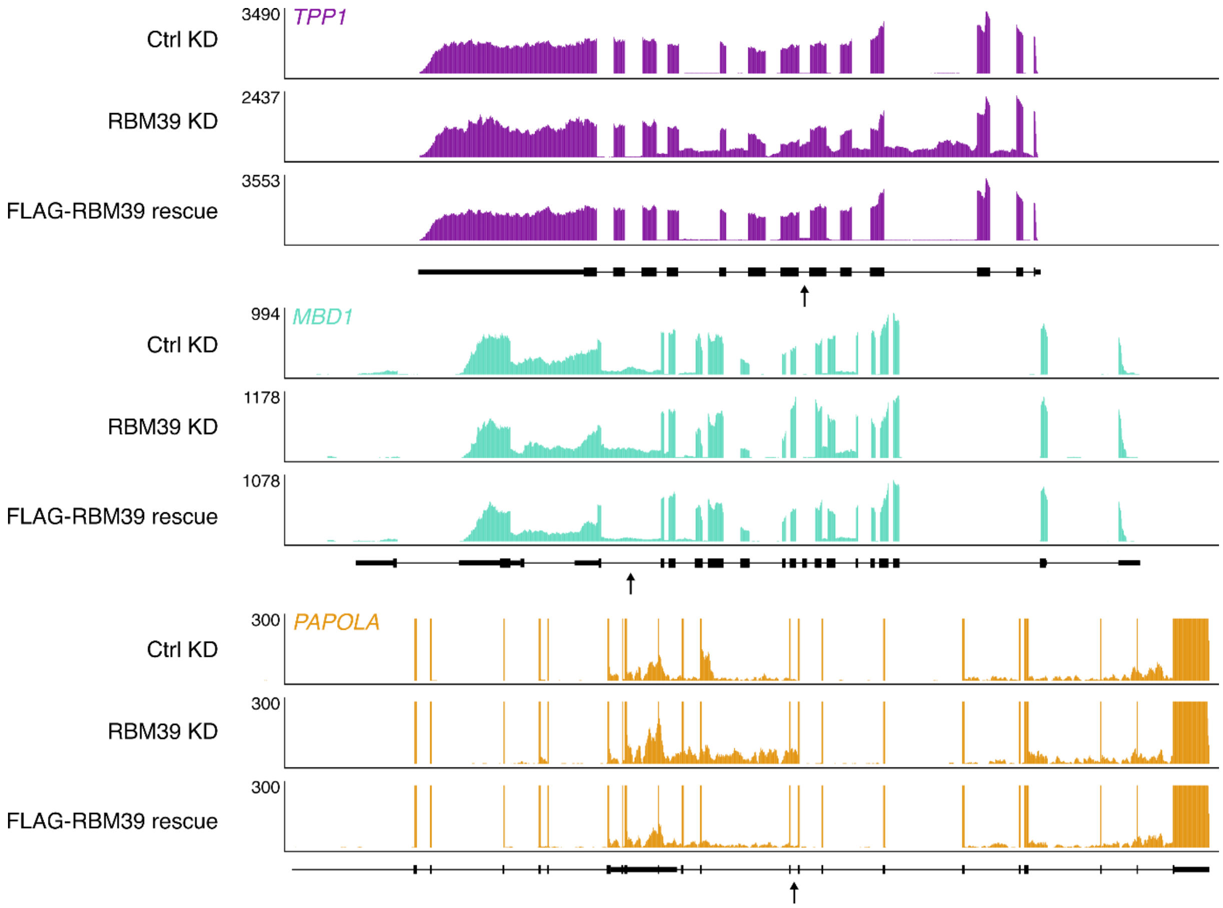


**Figure S2, related to Figure 2. RBM39-dependent intron retention events.** Sashimi plots showing RNA sequencing reads detected for three genes (*TPP1*, *MBD1* and *PAPOLA*) in conditions control (Ctrl KD), after RBM39 knock down (RBM39 KD) and after FLAG-RBM39 rescue (FLAG-RBM39 rescue). The intron retention events that were used for the validation of the RNA-seq data are marked by arrows. In the *TPP1* gene, several intron retention events are RBM39-dependant.

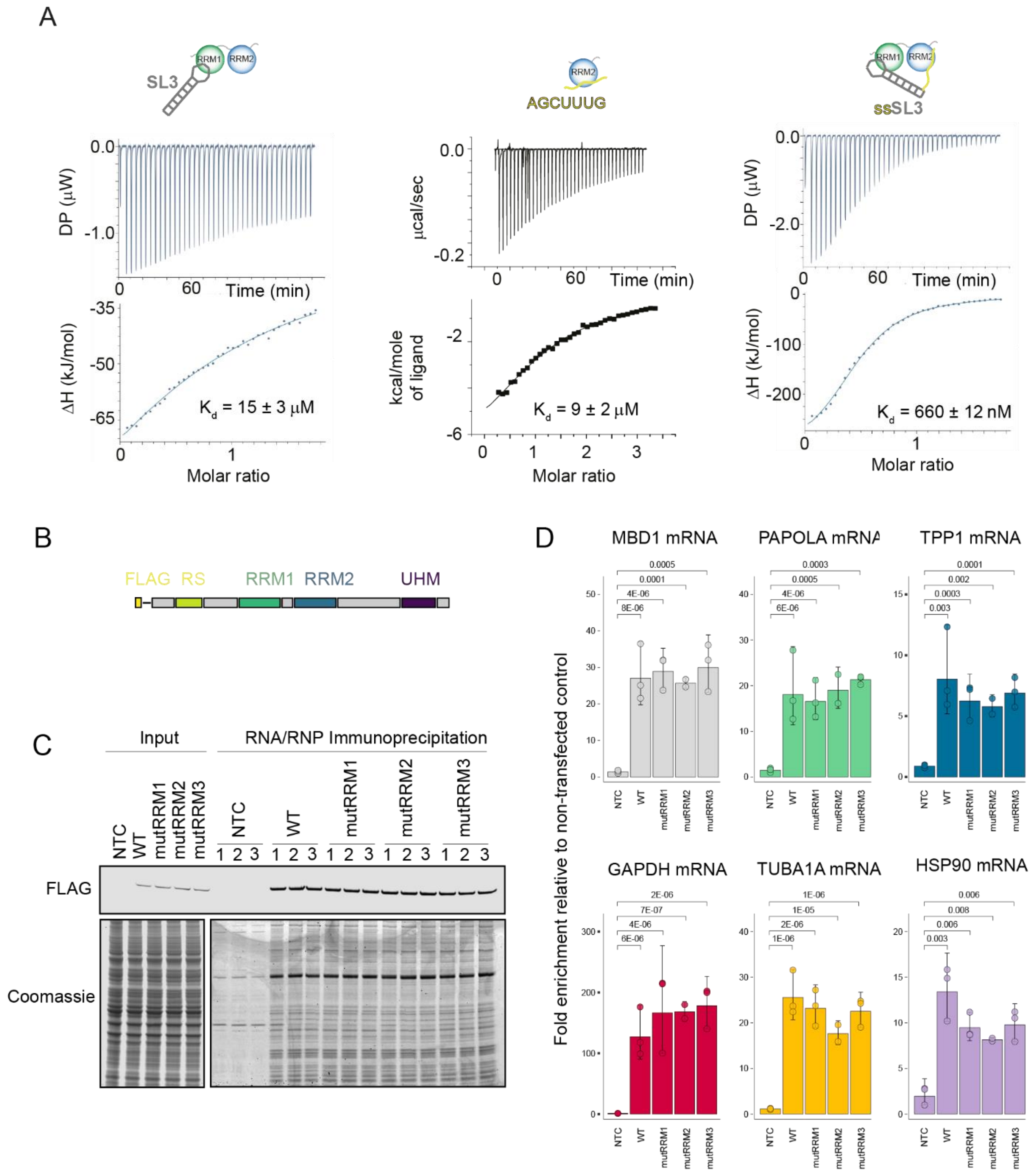


**Figure S3, related to Figure 3. ITC measurements and RNA immunoprecipitation with full-length RBM39.** A) From left to right, ITC titrations of RBM39 RRM12 with SL3, RRM2 with AGCUUUG and of RBM39 RRM12 and ssSL3. Average dissociation constants and errors are given on the plot (N=3). B) Schematic representation of FLAG-RBM39. C) On top, a western blot anti-FLAG reveals the presence of exogenous FLAG-RBM39 and its derivatives. At the bottom, Coomassie stained SDS-PAGE showing the protein composition of the input (left part) and the proteins that co-precipitate with FLAG-RBM39 and its derivatives. D) Bar plots showing the fold enrichment relative to non-transfected control for the MBD1, PAPOLA, TPP1, GAPDH, TUBA1A and HSP90 mRNAs immunoprecipitated with FLAG-RBM39 and its derivatives. P-values comparing each condition with the non-transfected control are given.

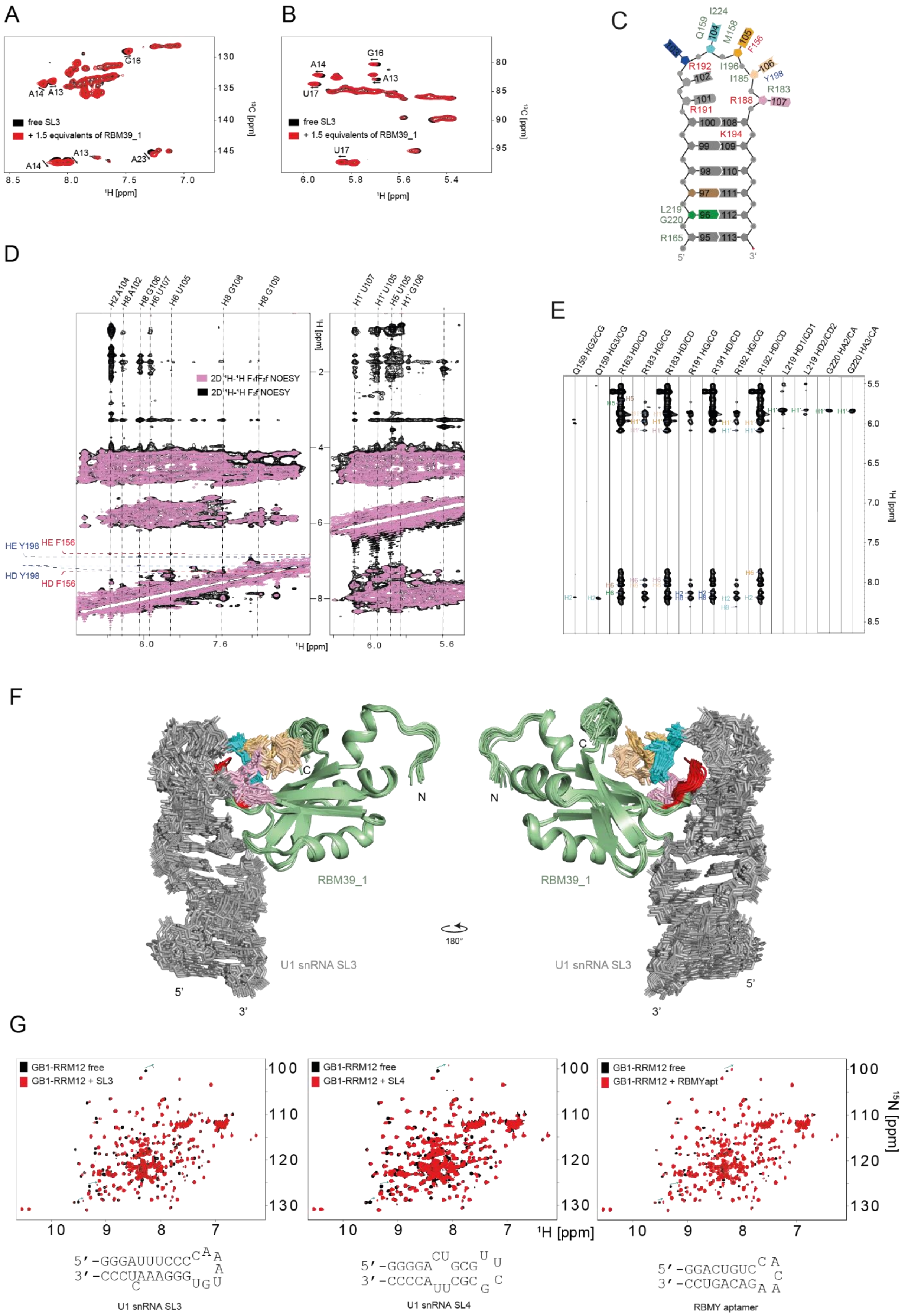

**Figure S4, related to Figure 4. RNA spectra, identification of intermolecular NOEs and solution structure of the RBM39 RRM1-U1 SL3 complex.** A) Overlay of the 2D $^{1}H$-$^{13}C_{aromatic}$ HSQC spectra of U1 SL3 before and after addition of unlabelled RBM39 RRM1. B) Overlay of the 2D $^{1}$H-$^{13}$C$_{sugar}$ HSQC spectra of U1 SL3 before and after addition of unlabelled RBM39 RRM1. C) Overlay of the 2D $^{1}$H-$^{1}$H F1fF2f NOESY and 2D $^{1}$H-$^{1}$H F2f NOESY spectra recorded with a sample of unlabelled U1 SL3 bound to $^{15}$N-$^{13}$C-labelled RBM39 RRM1. D) Selected strips of the 3D $^{13}$C-(F1 edited, F3 filtered) NOESY HSQC recorded with $^{13}$C-labelled RBM39 RRM1 and unlabelled SL3. E) Superimposition of the 20 solution structures of the RBM39 RRM1 – SL3 complex. G) Overlay of the 2D $^{1}$H-$^{15}$N HSQC spectra of GB1-RBM39-RRM12 before and after addition of U1 snRNA SL3, SL4 or of the RBMY aptamer. RNA sequences are given below the spectra.

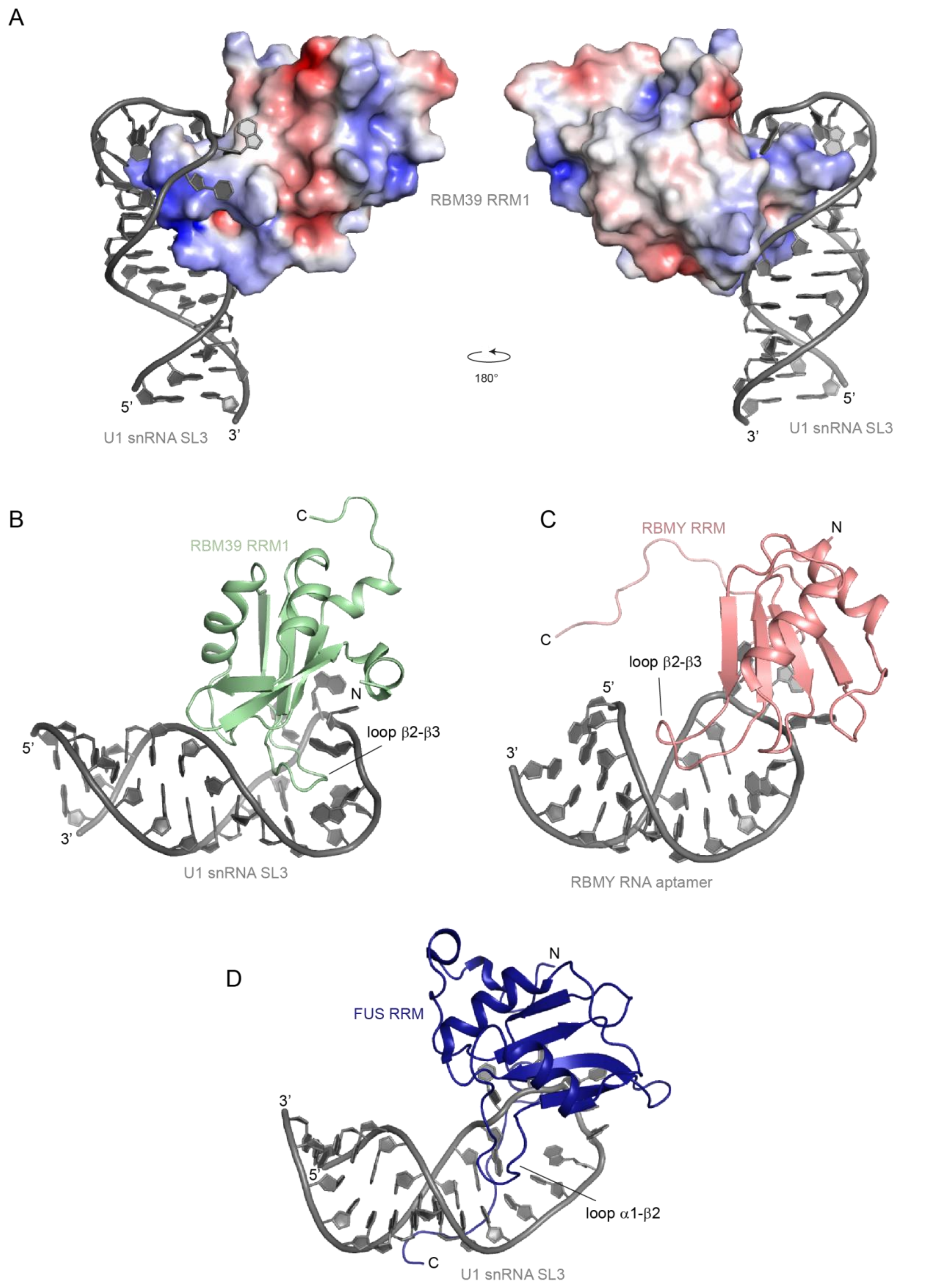


**Figure S5, related to Figure 4. Structural comparison between RBM39, RBMY and FUS RRM interactions with RNA stem loops.** A) Electrostatic potential surface of RBM39 RRM1 when bound to  U1 snRNA SL3. B) Cartoon representation of the lowest energy model of the NMR structure of RBM39 RRM1 bound to U1 snRNA SL3. C) Cartoon representation of the lowest energy model of the NMR structure of RBMY bound to RBMY RNA aptamer (PDB ID 2FY1). D) Cartoon representation of the lowest energy model of the NMR structure of FUS RRM bound to U1 snRNA SL3 (PDB ID 26SNJ).

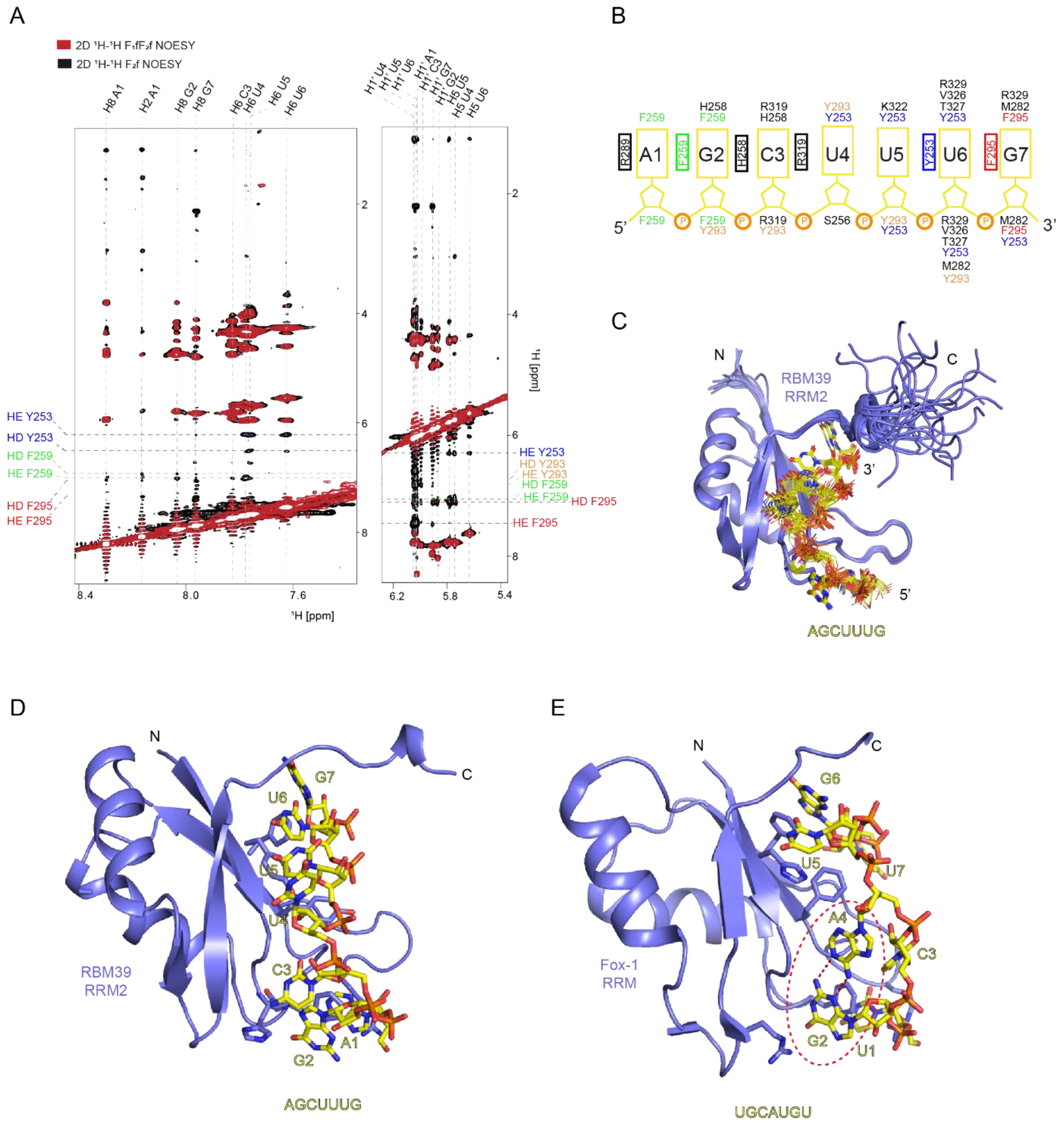


**Figure S6, related to Figure 5. Identification of intermolecular NOEs, solution structure of the RBM39 RRM2-AGCUUUG complex and comparison with Fox-1 RRM – RNA interaction.** A) Overlay of the 2D $^{1}$H-$^{1}$H F1fF2f NOESY and 2D $^{1}$H-$^{1}$H F2f NOESY spectra recorded with a sample of unlabelled AGCUUUG bound to $^{15}$N-$^{13}$C-labelled RBM39 RRM2. B) Schematic showing the intermolecular NOEs observed in the NMR data. C) Superimposition of the 20 solution structures of the RBM39 RRM2 – AGCUUUG complex. D) Cartoon representation of the lowest energy model of the NMR structure of the RBM39 RRM2 – AGCUUUG complex. E) Cartoon representation of the lowest energy model of the NMR structure of the Fox-1 RRM2 – UGCAUGU complex (PDB ID 2EER). The formation of the intermolecular base pair between A4 and G2 is highlighted by the red ellipse.

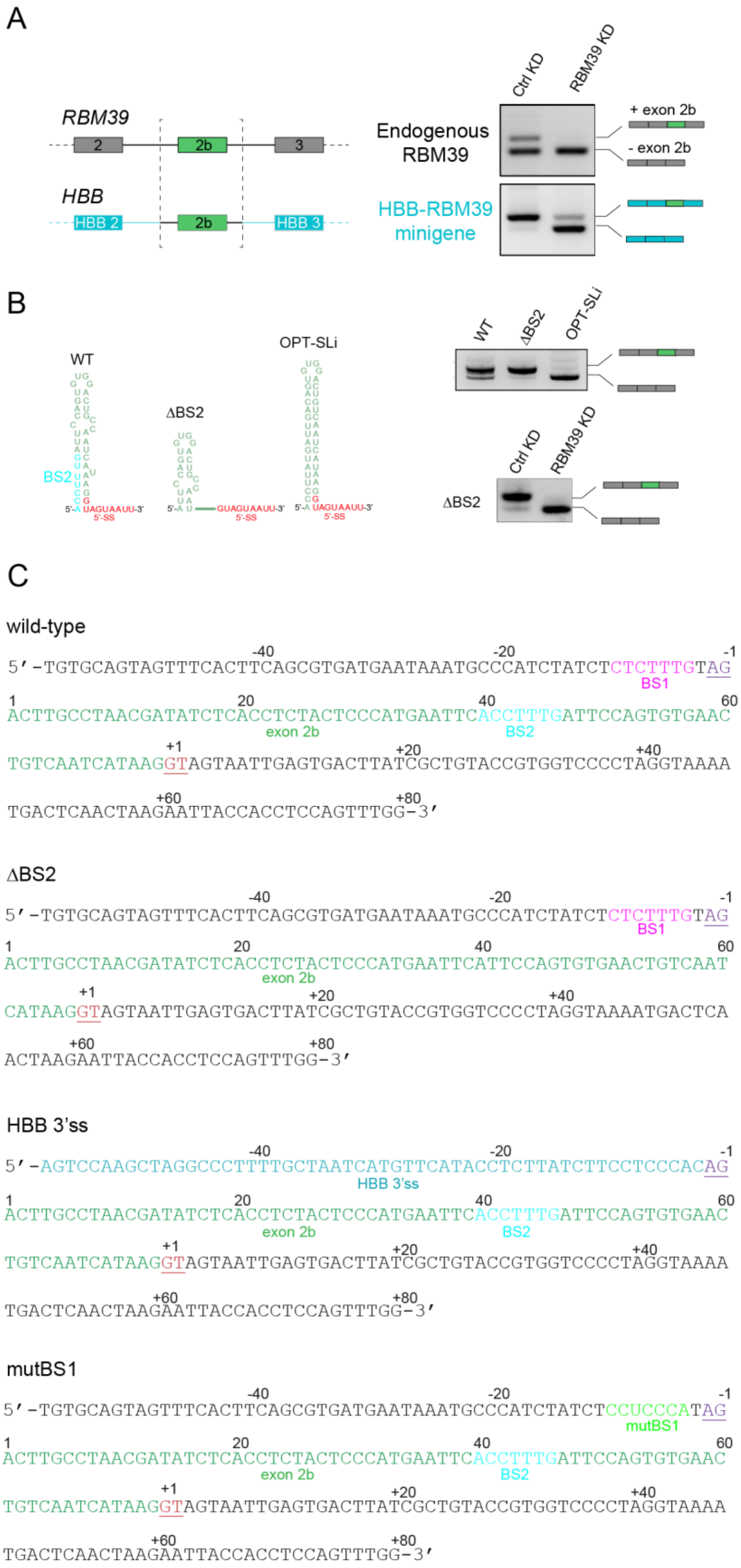


**Figure S7, related to Figure 7. Identification of the *cis* RNA element responsive for the RBM39 dependency of the poison exon inclusion.** A) Effect of the replacement of the RBM39 poison exon in the genomic context of the β-globin. The splicing of the poison exon was evaluated by RT-PCR in both context upon control KD or RBM39 KD. B) Evaluation of the effects of the predicted secondary on the splicing of the poison in the context of the RBM39 minigene. On the left, the different inhibitory stem loops are schematically represented. The mutation ΔBS2 releases the 5'-splice site from base pairing while the mutation OPT-SLi strengthens the secondary structure. Both mutations have an effect on the splicing of the poison exon: ΔBS2 favours poison exon inclusion while OPT-SLi induces constitutive skipping of the poison exon. However, the construct ΔBS2 is still sensitive to RBM39 KD, suggesting that BS2 is the not the *cis* RNA element responsible for the RBM39 dependency of poison exon inclusion. C) Sequences of the poison exon and its flanking regions used in Figure 7A-B.

**Table S1. NMR statistics of the RBM39 RRM1-U1SL3 and RBM39 RRM2-AGCUUUG complexes**

| | RRM1 – SL3 | RRM2 -AGCUUUG |
|---|---|---|
| **NMR restraints** | | |
| Distance restraints | 2,586 | 2,361 |
| Protein | | |
| Intramolecular | 2,200 | 2,214 |
| Intraresidual | 457 | 427 |
| Sequential (\|i - j\| =1) | 561 | 526 |
| Medium range (1 <\|i - j\| < 5) | 443 | 445 |
| Long range (\|i - j\| ≥ 5) | 730 | 784 |
| Hydrogen bonds[a] | 9 | 32 |
| RNA | | |
| Intramolecular | 319 | 58 |
| Intraresidual | 131 | 50 |
| Sequential (\|i - j\| =1) | 126 | 8 |
| Long range (\|i - j\| ≥ 5) | 52 | |
| Hydrogen bonds | 10 | |
| Complex | | |
| Intermolecular | 66 | 84 |
| Torsion angles[b] | | |
| Protein backbone | 144 | 150 |
| RNA angles | 138 | |
| **Energy statistics**[c] | | |
| Average distance constraint violations | | |
| 0.3-0.4 Å | 3.1 ± 1.8 | 6.4 ± 3.0 |
| > 0.4Å | 2.0 ± 1.2 | 2.6 ± 1.4 |
| Maximal (Å) | 0.51 ± 0.19 | 0.52 ± 0.19 |
| Average angle constraint violations | | |
| < 5 degrees | 29.8 ± 10.2 | 6.8 ± 2.5 |
| > 5 degrees | 2.5 ± 10.9 | 2.5 ± 0.9 |
| Maximal (degrees) | 6.09 ± 1.34 | 6.02 ± 3.96 |
| Deviation from ideal covalent geometry | | |
| Bond Length (Å) | 0.0038 ± 0.0013 | 0.0041 ± 0.0001 |
| Bond Angle (degrees) | 1.452 ± 0.484 | 1.404 ± 0.008 |
| Ramachandran Plot Statistics[c] | | |
| Residues in most favoured regions (%) | 87.0 ± 1.5 | 85.8 ± 1.4 |
| Residues in additionally allowed regions (%) | 11.7 ± 1.3 | 14.2 ± 1.2 |
| Residues in generously allowed regions (%) | 1.4 ± 0.9 | 0.0 ± 0.0 |
| Residues in disallowed regions (%) | 0.0 ± 0.0 | 0.0 ± 0.0 |
| RMSD to Mean Structure Statistics[d,e] | | |
| Protein | | |
| Backbone atoms | 0.40 ± 0.14[d] | 0.20 ± 0.04[e] |
| Heavy atoms | 0.86 ± 0.17[d] | 0.45 ± 0.06[e] |
| RNA | | |
| Backbone atoms | 0.79 ± 0.22[d] | 0.72 ± 0.16[e] |
| Heavy atoms | 0.82 ± 0.20[d] | 0.71 ± 0.15[e] |
| Complex | | |
| Backbone atoms | 0.68 ± 0.17[d] | 0.71 ± 0.16[e] |
| Heavy atoms | 0.92 ± 0.17[d] | 0.79 ± 0.15[e] |

RMSD, root-mean-square deviation. [a] Hydrogen bond constraints were identified from slow exchanging amide protons in $D_2O$ and imino protons in $H_2O$. [b] Torsion angle based TALOS+, sugar puckers based on homonuclear TOCSY, RNA backbone constraints in A form stem based on standard A form geometry. [c]Ramachandran plot, as defined by the program Procheck

(Laskowski *et al.*, 1996). [d]RMSD range 143-283 for the protein (chain ID: A) and 94-118 for the RNA (chain ID: B). [e]RMSD range 284-370 for the protein (chain ID: A) and 4-24 for the RNA (chain ID: B).

### Table S2. Oligonucleotides list

| Oligo ID | Oligo name | Sequence |
|---|---|---|
| | | **Cloning** |
| sc1 | RBM39 RRM1 fwd Nde1 | ATACATATGATTGATAATCTGACCCCG |
| sc2 | RBM39 RRM1 rev XhoI | CAGCTCGAGCGCAGCTGCGCGGTTTTTC |
| sc3 | RBM39 RRM2 fwd Nde1 | TATACATATGGCGGGTCCGATGCGTCTGTATG |
| sc4 | RBM39 RRM2 rev Nde1 | CCGCTCGAGTGCATCCGTGCGTTCGGTAACG |
| dj217 | RBM39 fwd BamH1 | GGGAGGATCCATGGCCGACGACATCG |
| dj218 | RBM39 rev Not1 | CTAGAATGCGGCCGCTCATCATCTCC |
| mdr835 | RBM39-dRRM3rev | ATATGCGGCCGCTCATCATTCGGTGTCCCAGCCCACTTCTTCC |
| dj282 | Sal1 RBM39 fwd | AAATGTCGACGCTGTCGCAGGG |
| dj283 | RBM39 ex4 Not1 rev | TTTGCGGCCGCTCAACGTTCTTCATGGCCGTTGG |
| dj734 | BamH1-RBM39-RRM1 fwd | AAAGGATCCATCGACAATCTGACCCCTGAG |
| dj735 | RBM39-RRM2-SV40NLS-STOP-Not1 rev | TTTGCGGCCGCTCATCACTTGTCCTCCACTTTGCGTTTCTTTTTG GGCTCGGTCACGTGGCCCAC |
| dj736 | RBM39 RRM12 linker fwd | CAACCTGCAGAAGGGAAGC |
| dj737 | RBM39 RRM12 linker rev | GCTTCCCTTCTGCAGGTTG |
| dj714 | Blg2-HBBe1_fwd | CCCAGATCTACTAGCAACCTCAAACAGAC |
| dj715 | HBBi2_rev-Hind3Sal1 | GTCGACGATCGTGCAAGCTTATGTACTAGGCAGACTGTG |
| dj716 | Hind3Sal1-HBBi2_fwd | AAGCTTGCACGATCGTCGACCCAAATCAGGGTAATTTTGC |
| dj717 | HBBe3_rev-Xba1 | GGGTCTAGATAGGCAGAATCCAGATGCTC |
| dj777 | Hind3-RBM39-i2-f | CGCAAGCTTCTTATCAGAGATGCATTACGTG |
| dj778 | Sal1-RBM39-i3-r | ACTGTCGACTCCCAAACCACTACATTCAC |
| dj779 | RBM39 deltaBS1 QC | CCTCTACTCCCATGAATTCTCCAGTGTGAACTGTCAAT |
| | | **Site directed mutagenesis** |
| mdr842 | QC RBM39 Y198A | GAAGCAAGGGAATCGCCGCCGTGGAATTCGTGGACG |
| mdr843 | QC RBM39 F156A | ACGCCCGGACCGTGGCCTGTATGCAGCTGG |
| mdr866 | QC RBM39 R192A | CGATTCCCTTGCTGGCTCTGCTGTTCCGGTCGCTGATC |
| mdr864 | QC RBM39 Y253A | GCAGGCTGCCCACGGCCAGTCTCATGGGTC |
| mdr865 | QC RBM39 F295A | GTCGCTGAAGGTGATGGCGCCGTAGCCCTTGCTG |
| mdr867 | QC RBM39 RRM2-2 | GTCCTCGGTGATGTTGGCGGCCAGGCTGCCCACGGCC |
| mdr848 | QC crypt SS RBM39 | GTCACACATCTCGCCGATAAGCAAACAGGGATTCACA CTGCTTTAGT |
| sc5 | QC RBM39 RRM2 F259A | GTATGTGGGCTCTCTGCATGCCAATATTACCGAAGATATG |
| sc6 | QC RBM39 RRM2 H258A | CTGTATGTGGGCTCTCTGGCTTTCAATATTACCGAAG |
| sc7 | QC RBM39 RRM2 R289A | GATGGATAGTGAAACCGGTGCTTCCAAAGGCTACGGTTTTATC |
| sc8 | QC RBM39 RRM2 H258AF259A | CTGTATGTGGGCTCTCTGGCTGCCAATATTACCGAAGATATG |
| | | **RT-PCR** |
| dj328 | RBM39 ex1 fwd | GCAGCAGCAGCAATCTCTTC |
| dj329 | RBM39 ex4 rev | TGGCACTGCTCAACTTGTTC |
| dj284 | RBM39 mini fwd | CGCTACCGGACTCAGATCTC |
| dj285 | RBM39 mini rev | ACGTTCTTCATGGCCGTTGG |
| | | **RT-qPCR** |
| dj296 | MBD1 spliced f | AACCCGCTTCCGAGATAC |
| dj297 | MBD1 spliced r | GCCTCCAGTCTACTGCTTTC |
| dj298 | MBD1 unspliced f | TTTCAGCCTGGGTGGAAC |

| | | |
|---|---|---|
| dj299 | MBD1 unspliced r | GGGTATGGGCCTTTACCTC |
| dj320 | PAPOLA spliced f | AGGACATCCTCACCTCATAAA |
| dj321 | PAPOLA spliced r | CCACTCAAAGCAAGACAGTTAG |
| dj306 | PAPOLA unspliced f | AGGACATCCTCACCTCATAAAG |
| dj307 | PAPOLA unspliced r | TGGGAAAGGTGAAGAGAAAC |
| dj312 | TPP spliced f | GCTCTGGCACCAGCAATAAC |
| dj313 | TPP spliced r | AACCACACGGGCTACTGATG |
| dj314 | TPP unspliced f | TCTGGCACCAGCAATAAC |
| dj315 | TPP unspliced r | ACTGCTCCAGGAACTATG |
| mdr256 | Sybr 5.8S rRNA f | GGTGGATCACTCGGCTCGT |
| mdr257 | Sybr 5.8S rRNA r | GCAAGTGCGTTCGAAGTGTC |
| sre231 | Sybr UPF1 f | CAAATCGACGTGGCGCTCTC |
| sre232 | Sybr UPF1 r | CCACGTTGCTTAGCTCTTCC |
| om368 | RP9P f | CAAGCGCCTGGAGTCCTTAA |
| om369 | RP9P r | AGGAGGTTTTTCATAACTCGTGATCT |
| mdr785 | Sybr RBM39 f | TGGCGGCAAGAATTCGAC |
| mdr786 | Sybr RBM39 r | GCTAGAGGCACTGAGCTAAC |
| **In vitro transcription** | | |
| sc9 | SL3 template | TCTAATACGACTCACTATAGGGATTTCCCCAAATGTGGGAAACTCCC |
| sc10 | SL4 template | TCTAATACGACTCACTATAGGGGACTGCGTTCGCGCTTTCCCC |
| sc11 | RBMY aptamer template | ATTCTAATACGACTCACTATAGGACTGTCCACAAGACAGTCG |
